# Environment-mediated vertical transmission fostered uncoupled phylogenetic relationships between longicorn beetles and their symbionts

**DOI:** 10.1101/2022.10.12.511864

**Authors:** Yasunori Sasakura, Nobuhisa Yuzawa, Junsuke Yamasako, Kazuki Mori, Takeo Horie, Masaru Nonaka

**Author notes:** Corresponding to: Yasunori Sasakura.

## Abstract

The Coleoptera Cerambycidae (longicorn beetles) use wood under different states (living healthy, freshly snapped, completely rot, etc.) in a species-specific manner for their larval diet. Larvae of some Cerambycidae groups have mycetomes, accessory organs associated with the midgut that harbor fungal symbiont cells. The symbionts are thought to improve nutrient conditions; however, this has yet to be shown experimentally. To deduce the evolutionary history of this symbiosis, we investigated the characteristics of the mycetomes in the larvae of longicorn beetles collected in Japan. Lepturinae, Necydalinae, and Spondylidinae are the only groups that possess mycetomes, and these three groups’ mycetomes and corresponding fungal cells exhibit different characteristics between the groups. However, the phylogenetic relationship of symbiont yeasts does not coincide with that of the corresponding longicorn beetle species, suggesting they have not co-speciated. The imperfect vertical transmission of symbiont yeasts from female to offspring is a mechanism that could accommodate the host-symbiont phylogenetic incongruence. Some Lepturinae species secondarily lost mycetomes. The loss is associated with their diet choice, suggesting that different conditions between feeding habits could have allowed species to discard this organ. We found that symbiont fungi encapsulated in the mycetomes are dispensable for larval growth if sufficient nutrients are given, suggesting that the role of symbiotic fungi could be compensated by the food larvae take. *Aegosoma sinicum* is a longicorn beetle classified to the subfamily Prioninae, which does not possess mycetomes. However, this species contains a restricted selection of yeast species in the larval gut, suggesting that the symbiosis between longicorn beetles and yeasts emerged before acquiring the mycetomes.

---

Longicorn beetle, fungi, yeast, symbiosis, mycetome, Lepturinae

## INTRODUCTION

Heterotrophic animals use symbiotic microorganisms to improve their uptake of nutrients. This kind of symbiosis is generally mutualism because host animals provide a space for microorganisms to proliferate comfortably with essential nutrients, while microorganisms support food digestion and supply nutrients (Lindsay et al., 2020; Modolon et al., 2020). Through symbiosis, animals have found ways to adjust to variable environments that are not always conducive to survival. Therefore, symbiotic systems are crucial for understanding the flourishing of animals on Earth.

The wood-eating groups in the coleopteran insects, which comprise the most prominent animal order, have significant relationships with fungi, particularly yeasts (Vega and Blackwell 2005; Gibson and Hunter 2010; Hulcr et al., 2017; Biedermann and Vega 2020). Many yeasts have been isolated from the frass, tunnel, and digestive tubes of coleopteran larvae (Suh et al., 2005; Endoh et al., 2011; Davis et al., 2015). In some groups, adults have a specific structure (the mycangium) that stores fungal cells and passes them to the next generation during oviposition (Tanahashi et al., 2010; Joseph et al., 2021). The fungi proliferate in the tunnel where larvae dwell and are incorporated into the larval digestive tube. These fungi are consumed not only as part of the larval diet (Toki et al., 2012) but are also thought to contribute to creating a suitable condition for the larvae and assisting in the digestion of wood. Indeed, many yeasts isolated from coleopteran larvae have cellulose-, cellobiose- and xylose-digesting activities (Davis et al., 2015).

The Cerambycidae family (longicorn beetles) is a widespread coleopteran group of wood-eating beetles. Larvae of longicorn beetles have a unique fungi-storing organ, the mycetome, near the anterior end of the midgut (Schomann, 1937; Nardon and Grenier, 1989; Grünwald et al., 2010). The mycetome consists of a mass of knob-shaped sacs that attach to the outer membrane (serosa) of the gut. Mycetome sacs are filled with nutritional particles, and fungal symbiont cells consisting primarily of yeasts dwell in the lumen or mycetocytes (Jurzitza, 1960). The fungal cells are constantly transferred to the gut lumen (Grünwald et al., 2010) and are thought to assist in the digestion of wood.

Although the mycetome seems essential for assisting the symbiosis with fungi, its function is still obscure. Indeed, the mycetome is not conserved among all longicorn beetle groups (Schomann, 1937). Therefore, this structure may have been acquired in some groups that require an additional benefit from symbiont fungi to increase their survival fitness, such as developing efficient digestion assisted by fungi. However, the advantages that symbiont fungi confer to longicorn beetles still need to be resolved. To deduce the evolutionary history of the symbiosis facilitated by the mycetome, we investigated the distribution of this structure in various species from different subfamilies of longicorn beetles in Japan. We inspected larvae from two families, six subfamilies, 61 genera, 73 species, and 289 individuals of longicorn beetles. Species of the subfamily Lepturinae are dominant among the species containing a mycetome; however, some Lepturinae species do not have this structure. We found that the loss of a mycetome is correlated with diet choice. The Japanese longicorn beetle groups/species that had or that lacked a mycetome in our analysis were in good accordance with those in a previous report of the presence/absence of symbiont yeasts in European species (Schomann, 1937). *Aegosoma sinicum* belongs to the subfamily Prioninae, whose members do not have a mycetome. We found that its larvae exhibit symbiosis with limited yeast species classified into the genus *Scheffersomyces*, suggesting that symbiosis with yeasts preceded the acquisition of mycetomes during the evolution of Cerambicydae.

## MATERIALS AND METHODS

### Animals

Most of the longicorn beetle larvae were collected from dead trees. We used cultured larvae to analyze the species whose larvae are difficult to capture from the wild. Their species names were identified by the cuticle structure on the thorax (Cherepanov, 1988; Koiwaya, 2017). When the thoracic cuticle structure could not identify species, genomic DNA was isolated from larval somatic tissues with a Wizard Genomic DNA Purification kit (Promega), followed by PCR amplification of 28S rDNA with ExTaq DNA polymerase hot start version (Takara-bio) and with the primers 5’-GAACTTTGAAGAGAGAGTT-3’ and 5’-CAGGCATAGTTCACCATCTT-3’. After the PCR fragments were purified with the Qiaquick Gel Extraction kit (Qiagen), the sequence was determined by the conventional Sanger method, which was used to identify species with the aid of 28S rDNA sequence isolated from adults (Accession number PP375410-PP375497). The larval midgut was surgically isolated with fine tweezers in phosphate-buffered saline (PBS) to observe mycetomes and collect gut materials. Frass was collected from the tunnel on wood with cotton swabs. Larvae in the tunnel were isolated together with frass to identify species.

### Identification of fungi

Mycetome contents, gut materials, frass or fluid from adult genitalia were spread on potato dextrose agar (PDA) plates (4-1126-05 Sani-Spec, Japan or 05709 Shimazu Diagnostics, Japan), including 15 μg/ml chloramphenicol and 50 μg/ml ampicillin. 1L PDA agar broth contains 4g potato extract, 20g glucose, and 15g agar. Eggs were immersed in PBS for a few minutes, and the supernatant was spread onto PDA plates. The plates were incubated for a few days at 25°C or longer at 4°C. DNA was extracted from yeast or yeast-like colonies by a microwave-based method (Izumitsu et al., 2012) and was subjected to amplification of the internal transcribed spacer (ITS) rDNA with ExTaq DNA polymerase hot start version (Takara-bio) and with the primers 5’-CTTGGTCATTTAGAGGAAGTAA-3’ and 5’-TCCGTAGGTGAACCTGCGG-3’ according to the previous method (White et al., 1990; Gardes and Bruns 1993). In this analysis, up to four colonies on a single plate were soaked together in 100 μl Tris-Cl buffer if the colonies were available, before DNA extraction. Sequences were determined from the amplified bands after purification with a Qiaquick Gel Extraction kit (Qiagen). When sequences could not be determined due to the low quality of the chromatographs, each colony was subjected to sequencing analysis separately. When the mycetome contents did not yield colonies because the symbiont fungi were unculturable, we isolated DNA from mycetomes with Isoplant II (Nippon-Gene) according to the manufacturer’s instructions, followed by PCR amplification and sequencing of rDNA of fungal cells.

The identified rDNA sequences of fungal cells were compared with the reported DNA sequences in the standard nucleotide collection (nr/nt) database using the blastn program (Altschul et al., 1997). The species names of the source organisms of the DNA sequence that exhibited the highest similarity in the blastn analysis are listed in the “Blastn Best Hit” column in Table S1. We used the source organism name to refer to the isolated fungal cells for further analyses. When an undescribed species (for example, *Scheffersomyces* sp.) exhibited the best-hit, its strain identifier (such as strain Y13) is referred. We have to note that it does not always guarantee the actual yeast species because in many cases, our rDNA sequences did not exhibit a perfect match to a known rDNA sequence (Table S1, “Blastn score” column).

The internal transcribed spacer (ITS) of *Scheffersomyces* sp. Y1052_SICYLG3 has one *Eco*RI restriction site. The yeasts isolated from adult genitalia of *Leptura ochraceofasciata* females were identified by the presence of the *Eco*RI site. After PCR amplification, the PCR fragments were digested by *Eco*RI, followed by agarose gel electrophoresis.

Mycetome-possessing Lepturinae and mycetome-lacking longicorn beetle groups are compared according to the number of larvae possessing symbiont yeasts, and the number of yeast species characterized in these groups. We performed two-tailed Chi-square test using R x64.4.1.2 and RStudio softwares (R Core Team 2021; RStudio Team 2020). In the latter comparison, we counted the “Blastn Best Hit” names of yeasts that formed colonies on PDA plates. *Encyclops olivaceus* and species in Necydalinae and Spondylidinae have unculturable symbiont fungi in their mycetomes. On the other hand, we collected only culturable yeasts from larvae in a mycetome-lacking group by streaking gut contents or frass onto the PDA plates. For unbiased comparisons, the identified unculturable fungi were excluded from these counts.

### Phylogenetic analyses

The sequences were aligned using the M-COFFEE program (https://tcoffee.crg.eu/apps/tcoffee/do:mcoffee; Notredame et al., 2000; Wallace et al., 2006). After removing the residues at the ends that seemed not to align properly due to the differences in the lengths between the sequences, maximum likelihood trees were constructed with the MEGA software version 11 (Tamura et al., 2021). Trees were tested with 1,000 bootstrap pseudoreplicates. The aligned nucleotides used for phylogenetic constructions are shown in Table S2.

The phylogenetic tree for Lepturinae beetles was constructed using 28S rDNA sequence. The phylogenetic trees of the yeasts isolated from longicorn beetle larvae were constructed as follows. Referring to the results of the blastn search, the ITS sequences of the yeast from the Lepturinae, Spondylidinae, and *Aegosoma sinicum* larvae were used for the phylogenetic tree construction. When a yeast species was detected from more than one larvae of the same longicorn beetle species, only one sequence was used for the analysis. The ITS sequences from *Hyphopichia rhagii* (isolated from *Rhagium femorale* and *R. japonicum*), *Candida wanchemiae* (isolated from *Anastrangalia scotodes*), and uncultured fungus clone OUT_F389 (isolated from *Megasemum quadricostulatum*) are considerably shorter than those from other yeasts and were removed from the analysis in view of their potential influence on alignment.

### Culture of Cerambycidae larvae

Our artificial diet was developed based on the work of Gardiner (1970). The artificial diet Insecta FII (Nosan Corporation, Japan) prevents the growth of fungi with the aid of anti-fungal chemicals that are not open by the company. Nine grams of Insecta FII, 6 g of dead *Symplocos coreana* powder, and 15 g of cellulose (ISC powder) were kneaded in the following liquid to create ISC paste: 30 or 45 ml of water, 30 or 45 ml of PDA dissolved in water, or 30 or 45 ml of 1x PDA suspended with one of two yeasts, *Candida vrieseae* or *Scheffersomyces* sp. Y1052_SICYLG3. These yeasts were cultured on PDA plates until colonies grew to cover the surface of the broth. The colonies were recovered with a cotton swab, and suspended in PDA liquid so that the liquid becomes cloudy. We did not quantify the suspended yeast cells. The yeasts were expected to improve larval growth; however, the effect was questionable and was not analyzed in detail. The ISC paste was spread in a polypropylene box and was autoclaved (20 min at 105°C) for sterilization. Eggs, hatched larvae or larvae collected from dead woods were placed on the autoclaved diet and were kept at room temperature until observation. For the introduction of selected yeast to larvae, eggs were sterilized for more than one day by placing them on sterilized ISC paste. The powder of PDA broth was suspended in distilled water and kept until agar powder sank. Then the supernatant was collected to obtain 1 x PDA liquid. A 70 mm x 30 mm x 2 mm rectangular wood plate of *Symplocos coreana* was sterilized by autoclaving, then applied with 1 x PDA liquid in which yeast cells were suspended by the same method as ISC paste. Sterilized eggs were cultured on the yeast-applied wood plate until hatching. Hatched larvae were collected and cultured in sterilized ISC paste for about one month until they became sufficiently large for the surgical isolation of mycetomes.

## RESULTS

### Distribution of mycetomes in Japanese Cerambycidae

To deduce the evolutionary history of the mycetome in longicorn beetles, we investigated the distribution of mycetomes in the Japanese species (Fig. 1). The majority of the cerambycid species we inspected representing the subfamilies Lepturinae, Necydalinae, and Spondylidinae, had mycetomes near the anterior end of the larval midgut (Fig. 1 and Table 1). There are group-specific characteristics in the shapes of mycetomes. The mycetomes of most Lepturinae species (Fig. 1B-C) and all Spondylidinae (Fig. 1D) adhered tightly to the serosa of the midgut. By contrast, the large mycetomes of Necydalinae species were well separated from the serosa with a connection through several thin tubes (Fig. 1E). The mycetomes of Lepturinae form a single circular ring surrounding the serosa (Fig. 1B). In contrast, those of Spondylidinae larvae form two narrower rings (Fig. 1D). The characteristics of the appearance of mycetomes in Lepturinae and Spondylidinae agree with those in the previous report (Buchner 1965).

**Fig. 1.**
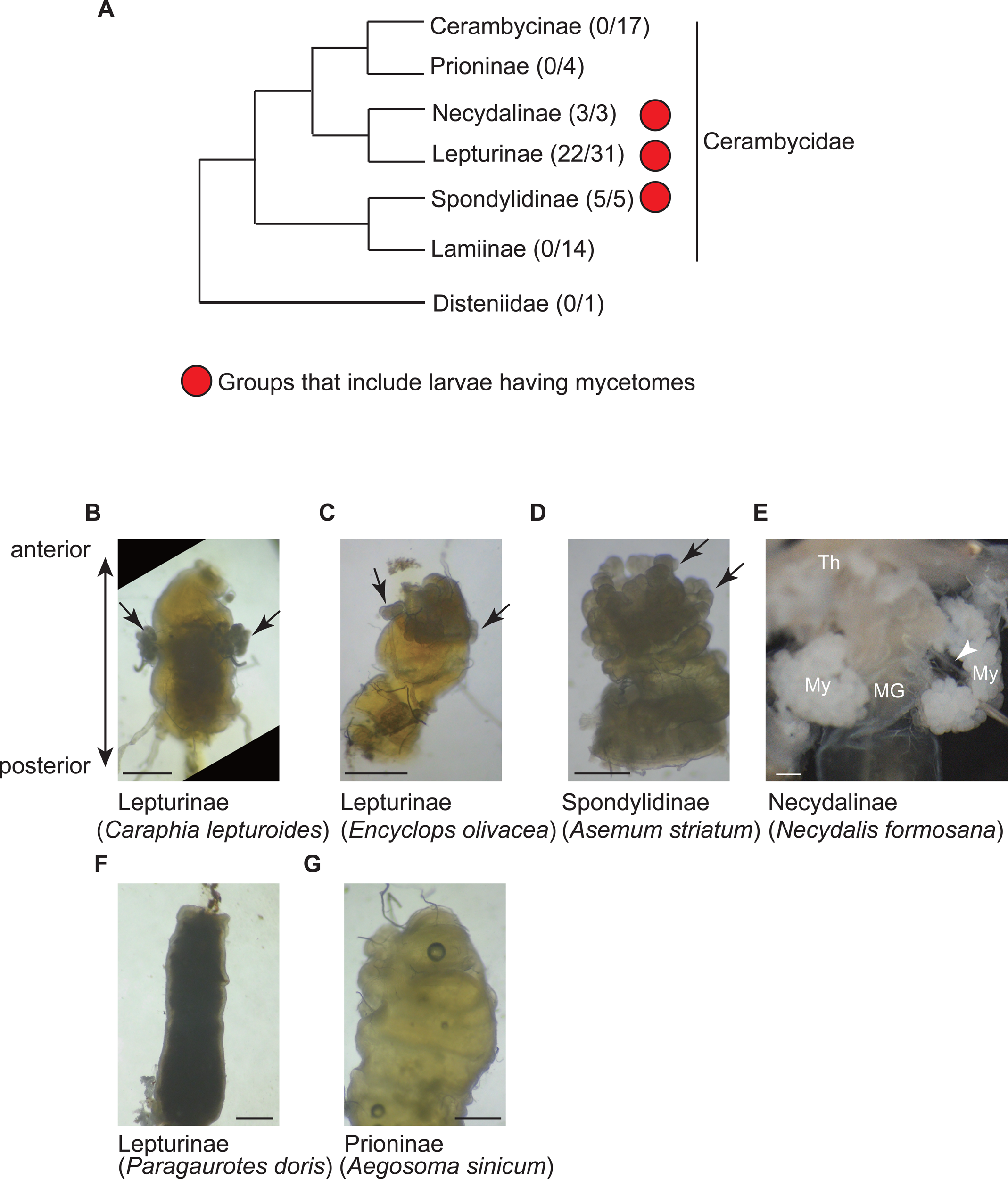
Detection of mycetomes in Japanese cerambycid larvae. **(A)** The phylogenetic relationships of Japanese longicorn beetle families/subfamilies. Red circles indicate groups including mycetome-possessing species. The number of species examined is shown in parentheses. This phylogenetic tree follows that of Hadded et al. (2018), although the branch length does not reflect phylogenetic distances. **(B)** Mycetomes (arrow) on the anterior midgut serosa of the Lepturinae *Caraphia lepturoides*. The anterior is toward the top. Scale bar, 500 μm. **(C)** Mycetomes of *Encyclops olivaceus*. Scale bar, 500 μm. **(D)** Mycetomes of the Spondylidinae *Asemum striatum*. Scale bar, 500 μm. The mycetomes of this group form two rings surrounding the serosa, which are shown by two arrows. **(E)** Giant mycetomes (My) of *Necydalis formosana*. Scale bar, 500 μm. MG, midgut; Th, thorax. The arrowhead indicates a thin tube that connects mycetomes with the serosa. **(F)** Mycetome-lacking midgut of the Lepturinae *Paragaurotes doris*. Scale bar, 500 μm. **(G)** Anterior midgut of the Prioninae *Aegosoma sinicum*. Scale bar, 500 μm. The mycetome is absent.

**Table 1.** Summary of symbiont characterization among longicorn beetle groups collected in Japan. The sum of yeast species isolated from Lepturinae having mycetome, Necydalinae and Spondylidinae, excluding duplicates between groups. The sum of yeast species isolated from Lepturinae without mycetome, Prioninae, Cerambycinae, Lamiinae, and Disteniidae, excluding duplicates between groups.

| Groups | Number of observed species | Number of observed larvae | N of yeast colony yielded larvae | N of larvae which had more than one yeast species | N of yeast species |
| --- | --- | --- | --- | --- | --- |
| Lepturinae having mycetome | 22 | 125 | 116 | 7 | 10 |
| Necydalinae | 3 | 12 | 0 | 0 | 0 |
| Spondylidinae | 5 | 19 | 3 | 1 | 8 |
| Lepturinae without mycetome | 9 | 38 | 13 | 5 | 15 |
| Prioninae | 4 | 25 | 20 | 2 | 11 |
| Cerambycinae | 17 | 38 | 15 | 3 | 13 |
| Lamiinae | 14 | 29 | 11 | 3 | 12 |
| Disteniidae | 1 | 5 | 4 | 0 | 4 |
| Mycetome present groups | 30 | 156 | 119 | 8 | 18 |
| Mycetome absent groups | 46 | 135 | 63 | 13 | 46 |

We found mycetomes in 22 out of 31 species of Lepturinae and all 3 of the examined Necydalinae species (Table 1). *Paragaurotes doris*, *Japanocorus caeruleipennis*, *Brachyta bifasciata,* and all *Pidonia* lacked mycetomes (Fig. 1F and Fig. 2). In these cases, the mycetomes were probably lost secondarily during speciation. Among the mycetome-lacking Lepturinae, larvae of *B. bifasciata* eat the roots of living *Paeonia* (Table S1, “Feeding habit” column). All of the other mycetome-lacking Lepturinae are inner bark-eaters at the larval stage. Inner bark-eating Lepturinae do not necessarily lack mycetomes; two *Rhagium* species are inner bark-eaters but have mycetomes. We found mycetomes in all examined Spondylidinae species (n=5; Table 1).

**Fig. 2.**
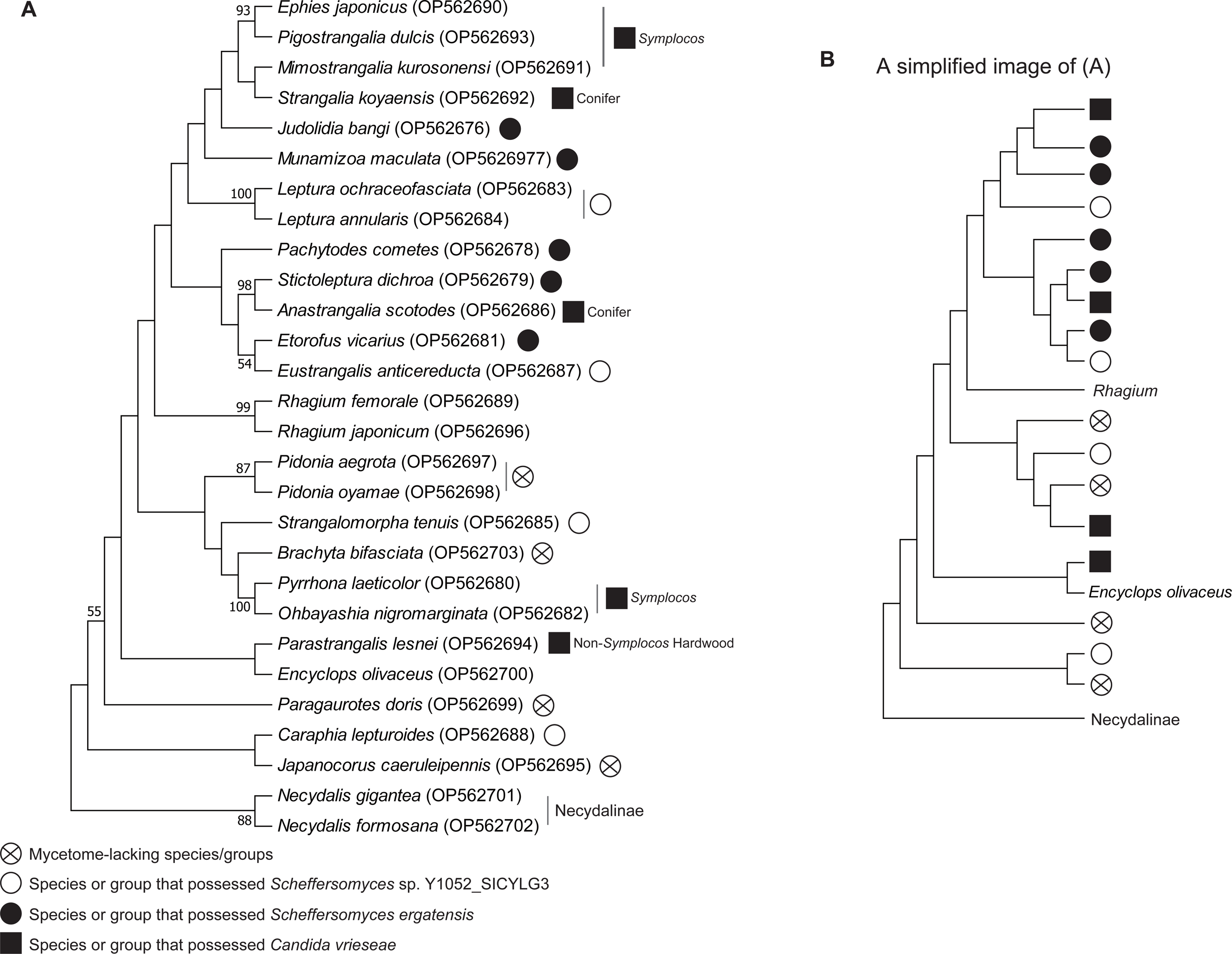
Distribution of mycetomes in the subfamilies Lepturinae and Necydalinae. **(A)** This phylogenetic tree was generated by the maximum likelihood method based on the alignment of 28S rDNA sequences. Crossed circles indicate mycetome-lacking species. Open circles indicate Lepturinae species that preferentially include the yeast *Scheffersomyces* sp. Y1052_SICYLG3 in their mycetomes. Black circles indicate Lepturinae species that preferentially include the yeast *Scheffersomyces ergatensis*. Squares indicate Lepturinae species that preferentially include the yeast *Candida vrieseae* with the information of their host trees. These three yeasts were frequently isolated from various species. Note that this tree does not include all species examined in this study for simplicity. The numbers beside branches indicate the percentage of times that a node was supported in 1,000 bootstrap pseudoreplication. The scores that are equal to or exceed 50% are shown. Species names are based on Danilevsky (2020). The accession numbers of the DNA sequences used for the phylogenetic tree construction are shown at the right side of the species names. **(B)** A simplified image of (A), highlighting the distribution of the symbionts and the loss of mycetomes.

### Characteristics of yeasts or yeast-like fungi in mycetomes

We found that mycetomes and fungi dwelling in them had group-specific characteristics. Each lobe of the Lepturinae mycetome contains tear-shaped fungal cells, a membranous structure, and particles much smaller than yeast cells (Fig. 3A and B, B’). The density of yeast cells in a mycetome is usually not high, and the cells do not fully occupy the lumen. When cultured on potato dextrose agar (PDA) plates, almost all of the fungi from the mycetomes of Lepturinae larvae formed colonies (Fig. 3C), and their rDNA sequences indicated that they were yeasts.

**Fig. 3.**
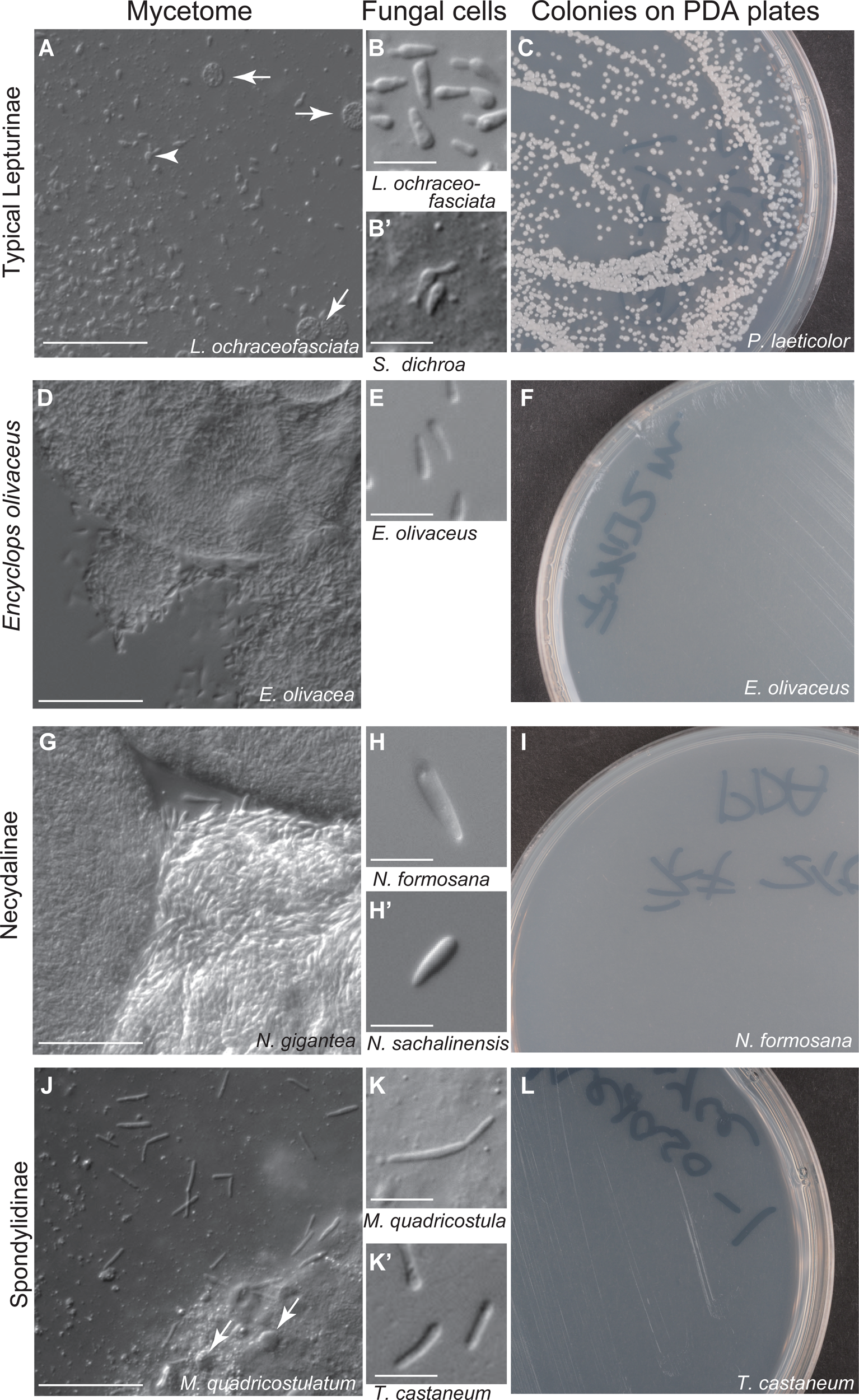
Fungal cells dwelling in the mycetomes. **(A-C)** Results for typical Lepturinae species. The names of the examined longicorn beetles are shown at the bottom of each panel. **(A)** Mycetome contents and yeast cells. Membranous structures are shown by arrows. An arrowhead indicates an example of symbiont yeast cells. Yeast cells are shown in higher-magnified images in **(B)** and **(B’)**. Scale bar, 50 μm in **(A)** and 10 μm in **(B)**. **(C)** Yeast colonies on a PDA plate. **(D-F)** The results for Lepturinae *Encyclops olivaceus*. In **(F)**, no colony was detected. **(G-I)** The results for Necydalinae species. The fungal cell in **(H’)** was isolated from a female oviduct. **(J-L)** The results for Spondylidinae species. Arrows indicate the membranous structures.

We classified yeasts isolated from the Lepturinae mycetome or gut by determining the sequence of the ITS site of mitochondria rDNA (Table S1). These rDNA sequences were used as queries of blastn searches to estimate the species of yeasts (Altschul et al., 1997). In the following section, we refer to the characterized yeasts as the species exhibiting the highest similarity to the rDNA sequences in the blastn analyses for simplicity.

In almost all cases, only a single yeast species was detected from the mycetome of a Lepturinae larva (87%, n=109/125; Table 1). However, seven larvae had more than one yeast species in the mycetome (rows 6, 8, 73-76, 107 in Table S1). Moreover, the rDNA sequence derived from the mycetomes of single larvae exhibited nucleotide variations in three cases (rows 10, 23, and 119 in Table S1). These results suggest that the yeast cells in mycetomes are not always clones. There are species-specificities between longicorn beetles and yeasts. For example, two Japanese *Rhagium* species had a yeast whose rDNA exhibited the best nucleotide sequence similarity to that of *Hyphopichia* (or *Candida*) *rhagii*, also found in a *Rhagium* species in Germany (Grünwald et al., 2010).

The phylogenetic analysis (Fig. 4) classified the yeasts isolated from Lepturinae mycetomes into eight species or unidentified species (*Scheffersomyces ergatensis*, *Scheffersomyces paraergatensis*, *Scheffersomyces coipomoensis*, *Scheffersomyces* sp. Y1052_SICYLG3, *Scheffersomyces* sp. Y6, *Scheffersomyces* sp. Y13, *Candida tenuis*, *Candida vrieseae*). Two indentified species (*Scheffersomyces* sp. Y6 and Y13) and *S. paraergatensis* are included into the branch of *S. ergatensis*. We did not include two species (*Hyphopicha rhagii* and *Candida wanchemiae*) in this analysis due to their short ITS sequences. In total, ten culturable yeast species were isolated from 22 mycetome-possessing Lepturinae species, suggesting that some yeast species were shared among different Lepturinae species (Table 1, Fig. 2, and Fig. 4). However, the Lepturinae species that share the same symbiont yeast are not always phylogenetically close to each other (Fig. 2). For example, the larvae of *Leptura ochraceofasciata* isolated from both hardwoods and conifers almost always had *Scheffersomyces* sp. Y1052_SICYLG3 (Table S1). This yeast was also found in the larvae of *Caraphia lepturoides*, although *L. ochracerofasciata* and *C. lepturoides* are phylogenetically distant. Larvae of *Pyrrhona laeticolor*, *Ohbayashia nigromarginata*, *Ephies japonicas*, *Pigostrangalia dulcis,* and *Mimostrangalia kurosonensis* had the yeast related to *Candida vrieseae* in their mycetomes. All of these Lepturinae species use the hardwood group Symplocaceae as their larval hosts. This was not due to the ubiquity of *C. vrieseae* in the dead Symplocaceae tree since larvae of the stag beetle *Aesalus asiaticus* isolated from a dead *Symplocos coreana* together with *Pyrrhona laeticolor* had a different yeast (*Scheffersomyces ergatensis*) in their digestive tube (Table S1, stag beetle). However, the relationships between yeast species and host plants of longicorn beetles are not expected to be rigid since *Anastrangalia scotodes* and *Strangalia koyaensis* use several conifers as hosts possessed *Candida vrieseae* (Table S1 and Fig. 4).

**Fig. 4.**
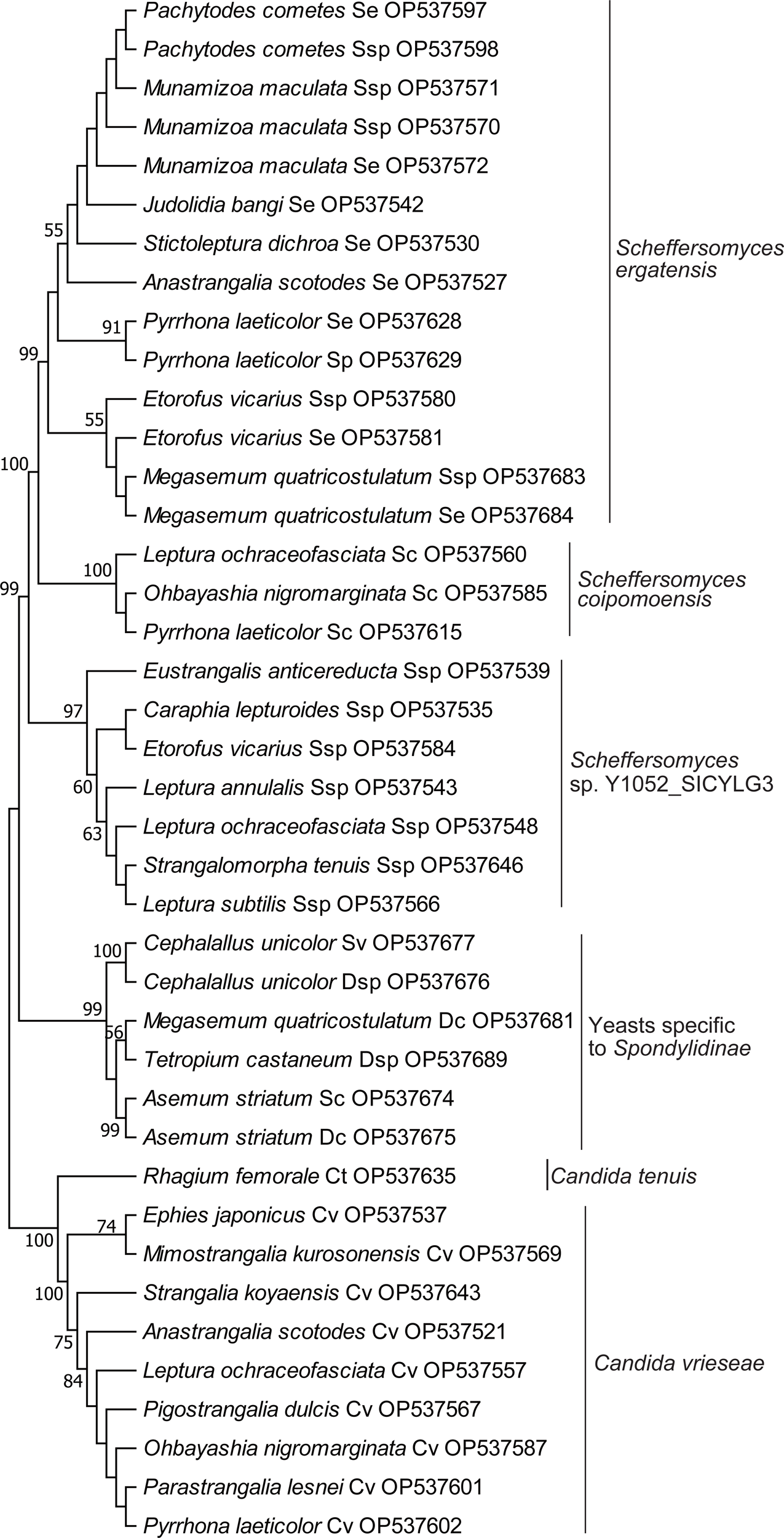
Phylogenetic relationships of yeasts isolated from the mycetome-possessing longicorn beetle larvae in this study. This phylogenetic tree was generated by the maximum likelihood method based on the alignment of ITS rDNA sequences. *Hyphopichia rhagii* (isolated from *Rhagium femorale* and *R. japonicum*), *Candida wanchemiae* (isolated from *Anastrangalia scotodes*), and uncultured fungus clone OUT_F389 (isolated from *Megasemum quadricostulatum*) were excluded from this tree because the yeast has much shorter ITS rDNA sequence than those of the other yeasts. The species names of yeasts are abbreviated. For example, “*Judolidia bangi* Se OP537542” indicates “*Scheffersomyces ergatensis* isolated from *Judolidia bangi*, and the accession number of the sequence used for the phylogenetic tree construction”. The number noted at a given branch indicates the percentage of times that a node was supported in 1,000 bootstrap pseudoreplications. The scores that are equal to or exceed 50% are shown.

The characteristics of the mycetome contents of the Lepturinae species *Encyclops olivaceus* and the three examined Necydalinae species were quite different from those of the other Lepturinae species. *Encyclops* and Necydalinae mycetomes were fully occupied by fungal cells, and we could not find a membranous structure (Fig. 3). The elongated fungal cells in these Necydalinae mycetomes were obviously larger than the yeast cells of Lepturinae, and the Necydalinae symbionts did not form a colony on PDA plates. The unculturable symbionts of *Encyclops* are similar in shape but smaller than those of Necydalinae species (Fig. 3E). All analyzed larvae of Necydalinae species are dependent on dead hardwood trees (Table S1). To further determine whether the symbiont characteristics are shared among Necydalinae species, we examined the symbiont of *N. sachalinensis*, which is dependent on the dead part of living conifers. Because its larvae were unavailable, we isolated fungal cells from the oviducts of *N. sachalinensis* females. The fungal cells had the same characteristics as those of the fungi isolated from the other Necydalinae larvae (Fig. 3H’ and Table S3).

We attempted to characterize the fungi from *Encyclops olivaceus* and Necydalinae mycetomes by directly isolating DNA from their mycetomes. The DNA fragments amplified from their mycetomes did not show sufficient similarity to those of any known fungi (Table S1). The DNA fragments from *Encyclops olivaceus* and *Necydalis sachalinensis* exhibited homology to the genome sequences of the other Coleopteran species (Table S1 and S3), remaining the possibility that these DNAs were not derived from the symbionts. Because Lepturinae symbiont yeasts yielded DNA bands specific to them by the same methods, these results confirm that the symbionts of the *Encyclops olivaceus* and Necydalinae species are distinct from yeasts.

The contents of Spondylidinae mycetomes were similar to those of typical Lepturinae in that it contained membranous structures, and the density of fungal cells was not high (Fig. 3J). The shape of Spondylidinae fungal cells was rod-like (Fig. 3K-K’). None of these fungi made a colony on PDA plates (Fig. 3L). Unlike Necydalinae, Spondylidinae fungal cells are classified into the yeast group *Debaryomyces*/*Schwanniomyces* (Table S1 and Fig. 4). These yeasts derived from Spondylidinae form a single clade in the phylogenetic tree of the yeasts isolated from mycetome-positive Lepturinae/Spondylidinae (Fig. 4). Therefore, there is broad group-specificity in the choice of symbiont fungi when Lepturinae, Necydalinae, and Spondylidinae are compared. However, when viewed from the species level, Lepturinae longicorn beetles’ phylogenetic relationships do not reflect in those of the symbiont yeasts (Fig. 2). This incongruence is our next target to address.

### Imperfect delivery of yeasts across generations

It has been suggested that the maternal parent provides the symbiont fungi in the larval mycetome (Schomann, 1937; Douglas, 1989). We characterized yeasts isolated from eight species of Lepturinae adult females. Although not perfectly matched, the adult females had yeast species isolated from the same species of larvae (Fig. 5; Table S3), confirming the vertical transfer of symbionts. We found that yeasts in the female genitalia could be easily excreted by gently pushing the oviduct. This suggests that yeasts are smeared around the eggs during oviposition, as previous studies described (Schomann, 1937; Douglas, 1989). Hatched larvae may incorporate yeasts with egg shells and nearby wood into their gut. Indeed, 1st instar larvae of *Ohbayashia nigromarginata* and *Necydalis formosana* a few days after hatching already had mycetome-like knobs at the anterior midgut and fungal cells in the structures (Fig. 6).

**Fig. 5.**
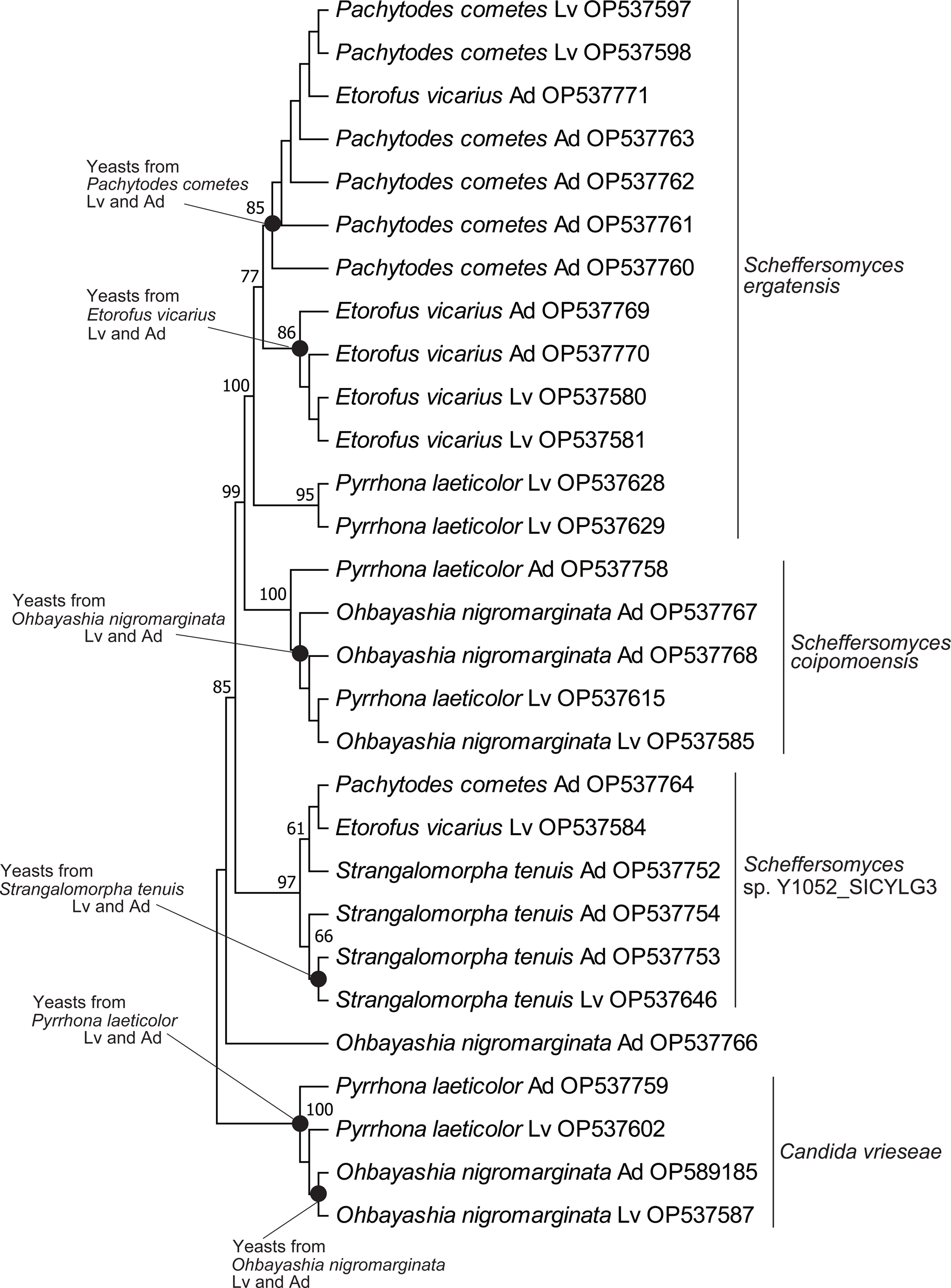
Phylogenetic relationships of yeasts isolated from larvae and adults of the Lepturinae beetles. This phylogenetic tree was generated by the maximum likelihood method based on the alignment of ITS rDNA sequences. The sequences of the yeasts isolated from the larvae are the same as the ones used in Fig. 4. Ad, yeast isolated from adult; Lv, yeast isolated from larva. The yeast species names are shown on the right side of the tree. For example, “*Pachytodes cometes* Lv OP537597” indicates “*Scheffersomyces ergatensis* isolated from a larva of *Pachytodes cometes*, and the accession number of the sequence used for the phylogenetic tree construction”. The clades that include the yeasts from both adults and larvae of the same Lepturinae species are marked with black dots. The number noted at a given branch indicates the percentage of times that a node was supported in 1,000 bootstrap pseudoreplications. The scores that are equal to or exceed 50% are shown.

**Fig. 6.**
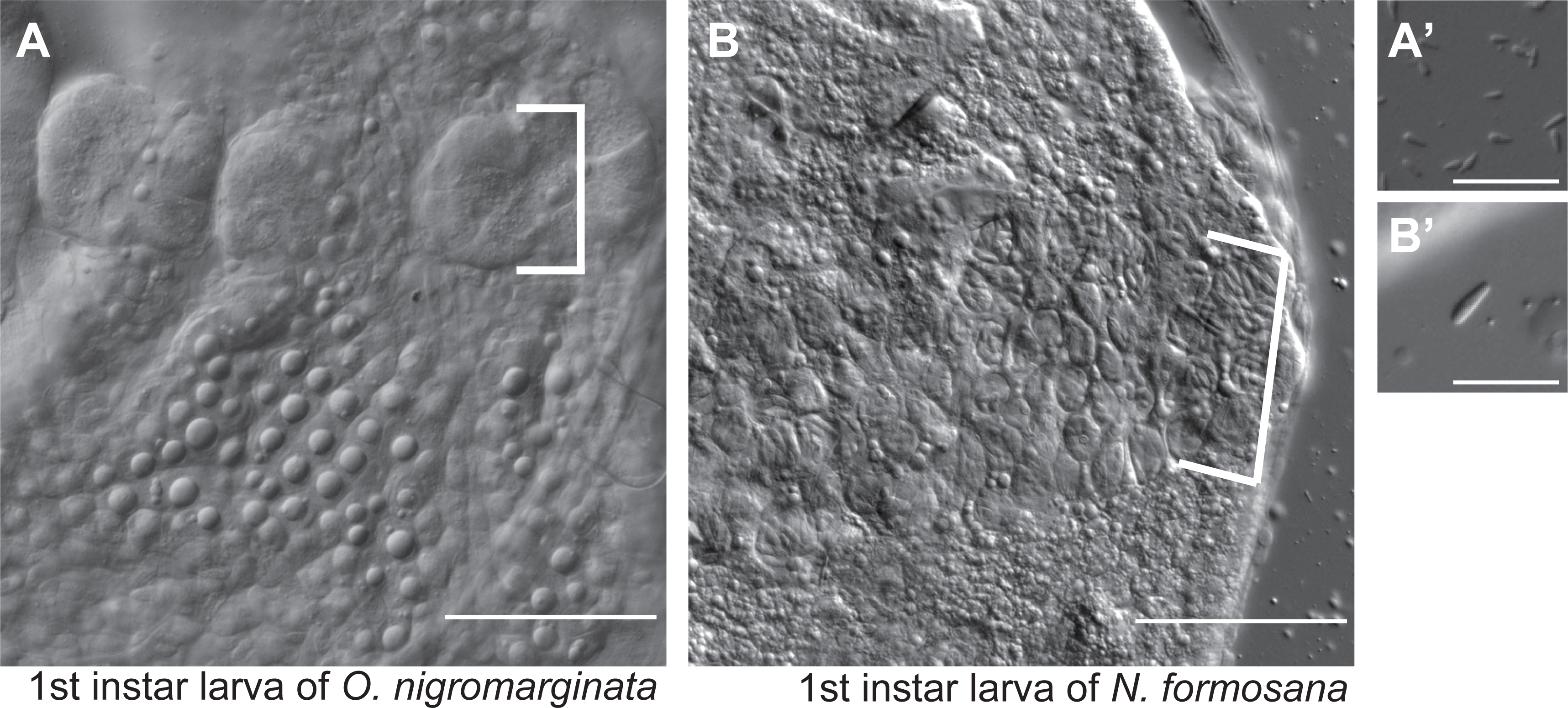
Fungal cells in the 1st instar larvae of Lepturinae/Necydalinae beetles. The observation was carried out over the first few days after hatching. **(A-A’)** The anterior midgut of *Ohbayashia nigromarginata* and yeast cells isolated from the gut. The bracket indicates mycetomes. Bar, 50 μm for A and 25 μm for A’. **(B-B’)** The anterior midgut of *Necydalis formosana* and a fungal cell isolated from the gut. Mycetome-like knobbed structures are shown by the bracket. Bar, 50 μm for B and 25 μm for B’.

This process of yeast delivery does not guarantee the accurate transmission of maternal yeasts to their descendants because larvae that accidentally move from the oviposited area before yeast uptake may have a chance to incorporate a different yeast species from the environment. For example, larvae of *L. ochraceofasciata* almost always have *Scheffersomyces* sp. Y1052_SICYLG3; however, we found a single larva with the yeast *Candida vrieseae* (Table S1). This individual was found in a dead tree of Symplocaceae. As we stated above, Lepturinae species that depend on this tree group usually have *Candida vrieseae*, and the larva of *L. ochraceofasciata* is suspected of having accidentally incorporated *Candida vrieseae* supplied by another Lepturinae species. We also found two larvae of *L. ochraceofasciata* and *P. laeticolor*, and five larvae of *Ohbayashia nigromarginata* that had the yeast *Scheffersomyces coipomoensis* (Table S1). These larvae were isolated from one dead tree of *Symplocos coreana* at Mt. Amagi. We isolated *Scheffersomyces coipomoensis* from yellowish dead trees of *Symplocos coreana* on the same mountain, suggesting that this yeast dwells in the environment and could have been accidentally incorporated into the larvae.

If Lepturinae larvae could incorporate yeasts from environment, it would be possible to generate yeast-free larvae or larvae having a yeast species different from that dominantly possessed by larvae in the wild populations, by removing maternally supplied yeasts. To show this, we developed a way to culture larvae under a condition chemically suppressing fungal growth. After oviposition, we placed eggs on an artificial diet containing anti-fungal chemicals for sterilization, so that the hatched larvae did not come into contact with external yeasts during and after birth. Larvae of *Leptura ochraceofasciata* and *Stictoleptura dichroa* grew well under this condition (Fig. 7A). They had brownish mycetomes near the anterior end of the midgut (Fig. 7B and D). The position of their mycetomes was identical to those of larvae from the wild. Microscopic observations showed that these brown mycetomes did not include yeasts (Fig. 7C and E), which was confirmed by culturing mycetome contents on PDA plates (Fig. 7C, inset), suggesting that the yeasts needed to be supplied externally for larvae to acquire yeasts in their mycetomes. This experiment also indicates that mycetomes can develop without yeasts.

**Fig. 7.**
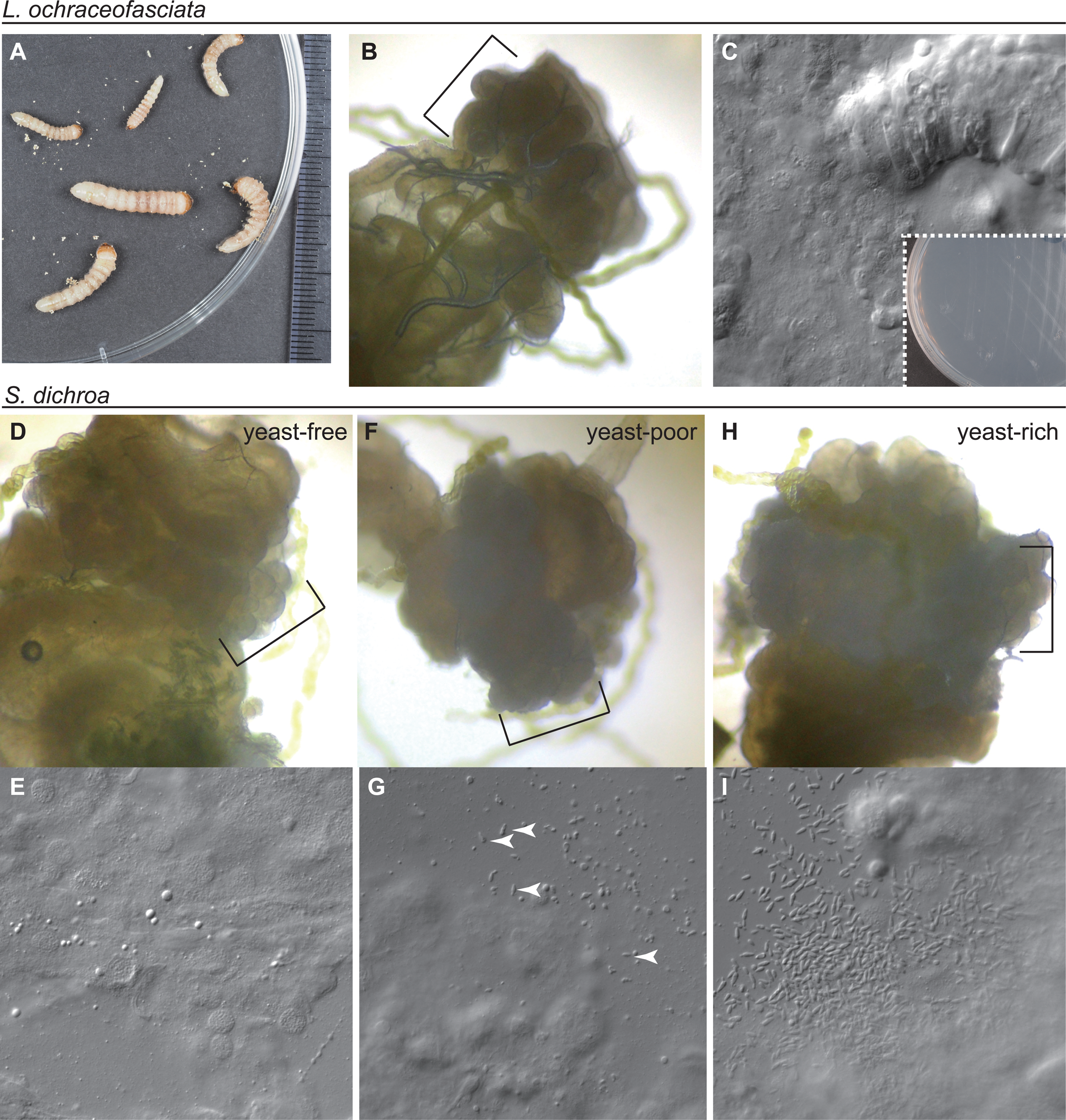
Yeast-free mycetomes in Lepturinae larvae. **(A)** *Leptura ochraceofasciata* larvae cultured with an artificial diet. The images were obtained approximately one month after hatching. **(B)** Yeast-free mycetomes (knobs shown by the bracket). **(C)** A magnified image of the contents of yeast-free mycetomes. The inset shows the results of culturing of the contents on a PDA plate. No colony appeared. **(D-I)** The results for *Stictoleptura dichroa*. **(D-E)** Yeast-free mycetomes. **(F-G)** Yeast-poor mycetomes. Note that the color of the mycetomes is somewhat whitish. Four examples of yeast cells are shown by arrowheads. The yeast *Meyerozyma caribbica* was identified from the mycetomes. **(H-I)** Yeast-rich mycetomes. The yeast *Scheffersomyces ergatensis* was identified from these mycetomes.

In the above culture experiments, some of the larvae had yeasts not found in the mycetomes of larvae isolated from wild populations. In the experiment with *L. ochraceofasciata*, we discovered that a few larvae (12.5%, n=2/16) had *Meyerozyma caribbica*. *Meyerozyma* is an indigenous yeast group arising in our laboratory. In the experiment using *S. dichroa*, a portion of eggs hatched on the wood onto which their mothers had laid eggs before sterilization. We cultured the pre-hatched eggs and 1st instar larvae in the same container. After one month, we observed that the larvae were divided into three groups: larvae with yeast-free mycetomes (n=5; Fig. 7D and E), larvae with mycetomes that included yeast cells with a low density (n=4; Fig. 7F and G), and larvae with relatively yeast-rich mycetomes (n=9; Fig. 7H and I). The yeast species isolated from the yeast-poor mycetomes differed from those isolated from yeast-rich mycetomes: the former was *Meyerozyma caribbica,* while the latter was *Scheffersomyces ergatensis*. *S. ergatensis* is the default species of this longicorn beetle in the wild (Table S1). We suspected that the larvae with *S. ergatensis* hatched on the egg-laid wood and had a chance to incorporate maternally gifted yeast.

To investigate the above hypothesis about environmental yeast acquisition, we next tested whether we could deliver a specific yeast species into mycetomes. In this experiment, sterilized eggs were placed on a wood plate to which a yeast species was applied. After birth, we transferred the larvae to a plate with the artificial diet to avoid further yeast incorporation during growth until the larvae reached a sufficient size to isolate mycetomes. We examined 33 larvae, and 22 had the given yeast species (Table S4). These yeasts were never detected in the control groups to which we did not provide the yeasts (0%, n=50), suggesting that this experiment successfully replaced the symbiont yeast with the arbitrary yeast. We concluded from these results that Lepturinae longicorn beetles acquire yeasts externally after hatching. Although the yeasts are usually maternally supplied, a different yeast could be obtained from the environment.

### Lepturinae mycetomes protect symbionts from anti-fungal chemicals in food

During the culture of longicorn beetle larvae with the artificial diet, we found that the mycetomes of Lepturinae species protect the symbionts from the anti-fungal chemicals in the artificial diet. A larva of *Etorofus vicaria* and *Leptura ochraceofasciata* maintained their symbiont yeasts in the mycetomes even after culturing for about one year (Fig. 8A and Table S5). By contrast, symbionts of *Necydalis formosana* larvae were eliminated from their mycetomes (Fig. 8B) even though they increased their weight during the culture (Table S5). These results suggest that the mycetome of this species is weaker in protecting symbionts from chemicals taken with the food.

**Fig. 8.**
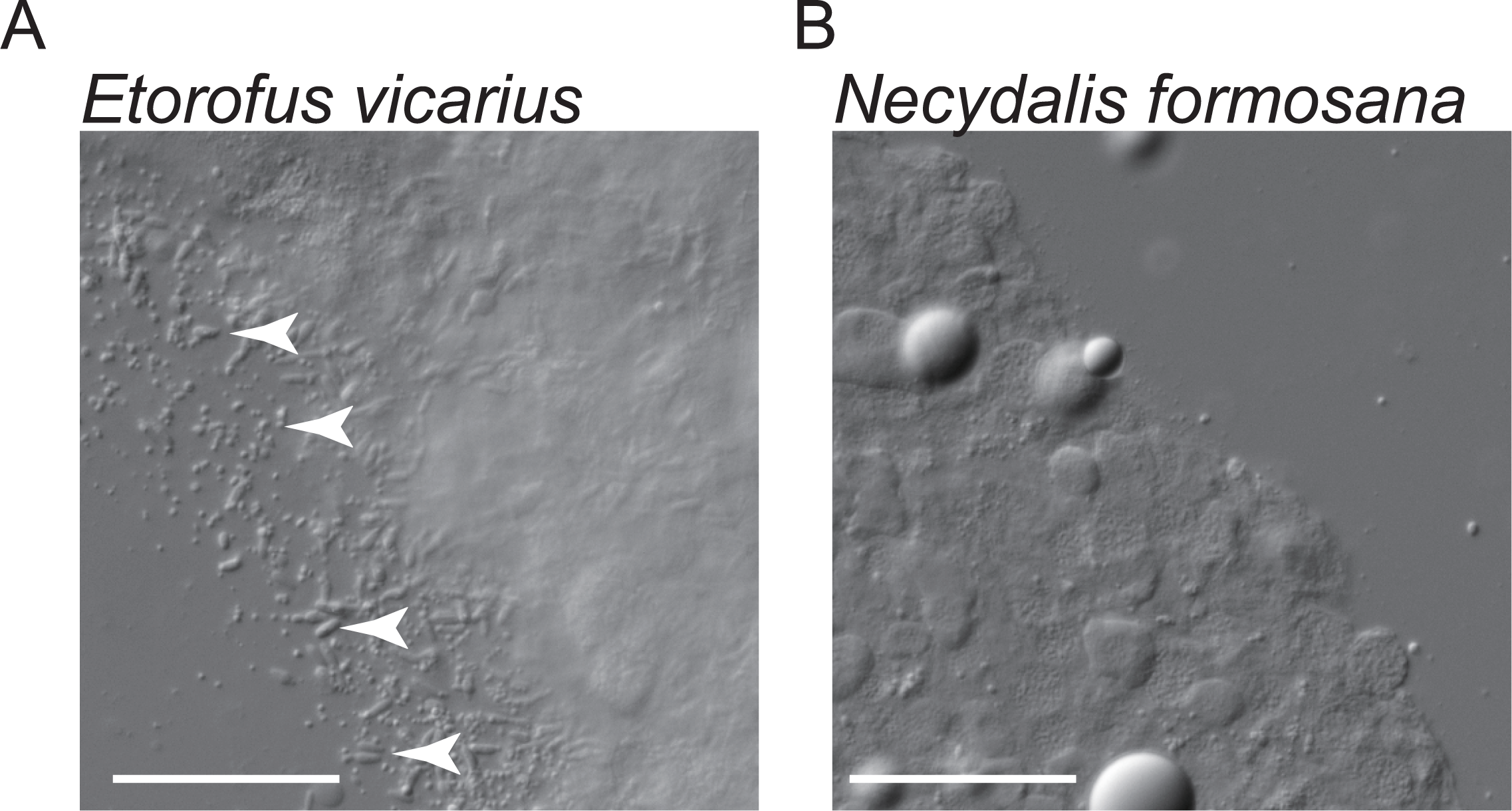
The presence of symbionts in the mycetomes after long-time culture on the artificial diet with anti-fungal chemicals. **(A)** *Etorofus vicarius* mycetome. Arrowheads are examples of symbiont yeasts. Bar, 100 μm. **(B)** *Necydalis formosana* mycetome. Bar, 100 μm.

### Yeast transmission between generations in the longicorn beetles without mycetomes

There is an association between mycetome-possessing Lepturinae longicorn beetles and yeast species. Our next question is how their symbiosis started, and notably, how mycetomes contributed to the evolution of the symbiosis. To address these questions, we investigated yeasts in the digestive tubes and frass of mycetome-lacking longicorn beetles. We observed 135 larvae of mycetome-lacking longicorn beetles (including Lepturinae) corresponding to 46 species from five families/subfamilies (Table 1). Yeasts were detected in 46% (n=63/135) of larvae. This score is significantly low compared to that of mycetome having Lepturinae (92%, n=116/125, *p*<0.001), suggesting that mycetomes have a role in stabilizing symbiosis with yeasts. We isolated in total 45 culturable yeast species from 46 mycetome-lacking longicorn beetle species (Table 1). This score is two times more than mycetome-having Lepturinae species (10 out of 21 excluding *Encyclops olivaceus* whose symbiont is unculturable; *p*<0.001), suggesting that yeasts habiting in the mycetome-lacking species are more variable than mycetome-possessing species. These wobbles could be explained if mycetome-lacking longicorn beetles do not usually possess a way to limit the species of symbiont yeasts, such as a delivery system between generations.

We found an exceptional mycetome-lacking species that could transmit specific yeasts between generations. *Aegosoma sinicum* is a kind of Prioninae that does not have mycetomes (Fig. 1G). Two yeast groups belonging to the genus *Scheffersomyces* were repeatedly isolated from larvae of *A. sinicum* collected in different locations and from different host trees (Table S1 and Fig. 9), suggesting a strong correlation between *A. sinicum* and the yeasts. Individual *A. sinicum* larvae exclusively possessed either of the two *Scheffersomyces* groups.

**Fig. 9.**
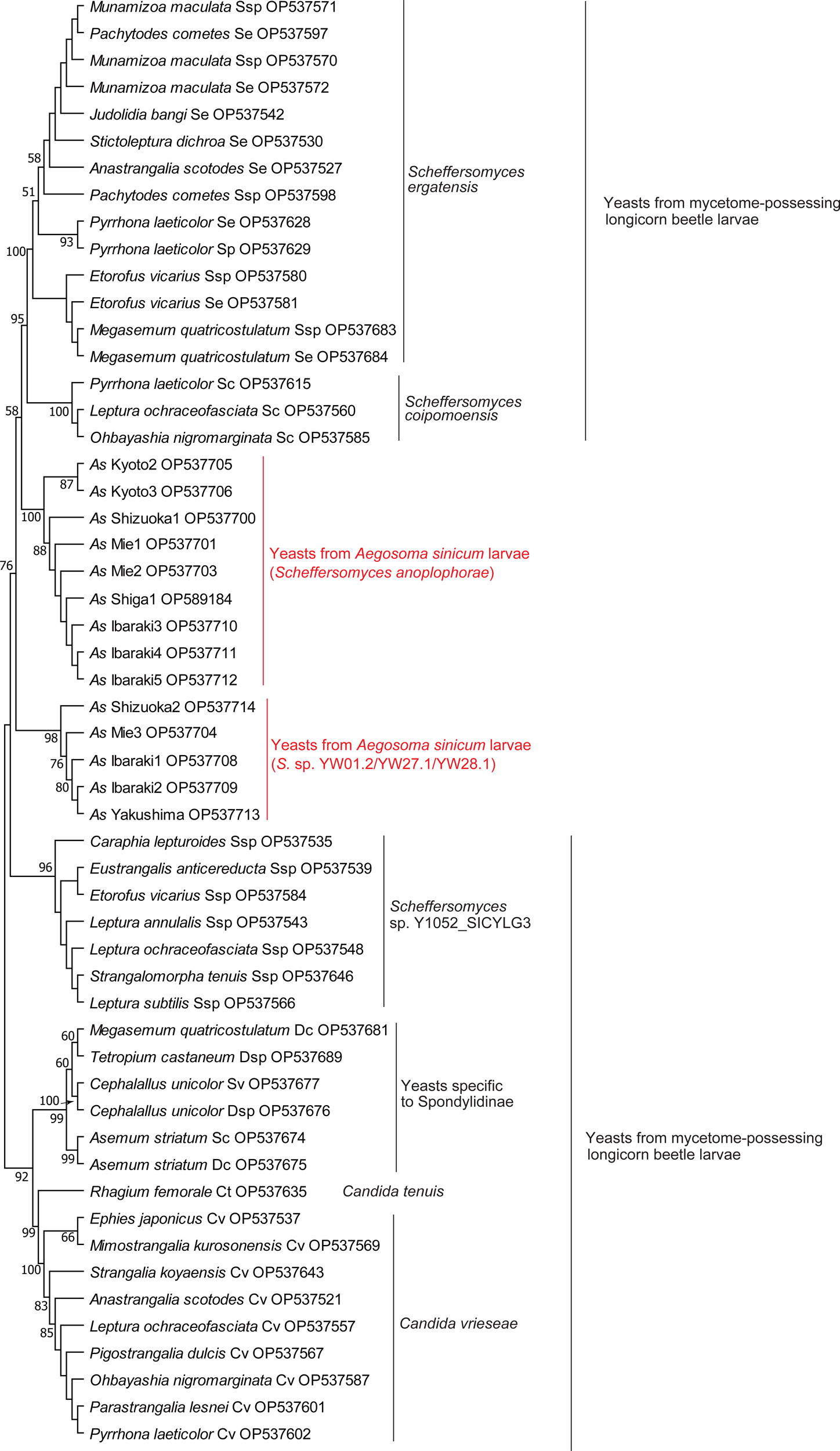
*Aegosoma sinicum* larvae have a symbiotic relationship with the two yeast groups. This phylogenetic tree was generated by the maximum likelihood method based on the alignment of ITS rDNA sequences. The yeasts from mycetome-possessing longicorn beetles used in Fig. 4 are also shown in this phylogenetic tree to investigate the phylogenetic relationships between mycetome-dependent and *Aegosoma* yeasts. The yeasts isolated from larvae of *Aegosoma sinicum* are indicated by the sampling location and identification numbers corresponding to those in Table S1. The number noted at a given branch indicates the percentage of times that a node was supported in 1,000 bootstrap pseudoreplications. The scores that are equal to or exceed 50% are shown.

Two stand out among the possible explanations for the relationship between *A. sinicum* and *Scheffersomyces*. The first is that *A. sinicum* females might lay eggs on the host trees in which the yeasts habit. The second possibility is that the yeasts were transferred from adult females to their progeny. We isolated the colonies of the same *Scheffersomyces* yeasts from the genitalia of females and the egg surfaces of *A. sinicum* (Table S2). Therefore, it is likely that *A. sinicum* vertically transfers yeasts between generations without mycetomes, although we cannot rule out the first possibility.

## DISCUSSION

Many dead wood-eating beetles depend on yeasts or related fungi at their larval stage (Davis, 2015). Although the role of this symbiosis has yet to be fully understood, its suspected function is to increase efficiency in the nutrient absorption from dead wood that is less nutritious than living plants. Larvae of longicorn beetles, which form one of the most diversified groups in the Coleoptera, evolved a characteristic fungi-storing organ, the mycetome, at the midgut (Schomann, 1936; Buchner, 1965). Among six subfamilies of Cerambycidae in Japan, the species in Lepturinae, Necydalinae, and Spondylidinae have mycetomes. The Japanese species in Disteniidae, which is a sister group of Cerambycidae, does not have the mycetome, confirming the previous report (Svacha and Lawrence 2014). Svacha and Lawrence (2014) also reported that the larvae in the families Vesperidae and Oxypeltidae do not have mycetomes. Because mycetome-absent families are dominant in longicorn beetles, mycetome was likely to have evolved in the subfamilies of Cerambycidae. If this is true, the mycetome emerged at least twice in Spondylidinae and Lepturinae/Necydalinae according to the phylogenetic tree of the Cerambycidae subfamilies (Fig. 1; Haddad et al., 2018; Nie et al., 2021). The emergence of the mycetome in Lepturinae and Necydalinae is likely to have been the single event at the root of these groups. This is supported by the recent phylogenetic study suggesting that Necydalinae is regarded to be included in Lepturinae (Nie et al., 2021). However, we cannot rule out the possibility that these mycetomes were evolved independently because Necydalinae mycetomes have characteristics that are not seen in Lepturinae mycetomes.

The symbionts of Necydalinae species and that of *Encyclops olivaceus* are unculturable and fully occupy the lumens of the mycetomes. The shared characteristics of the mycetomes and symbionts among *Encyclops* and Necydalinae species suggest that they are phylogenetically related. The previous study showed that the Lepturinae species *Oxymirus cursor* has gigantic mycetomes fully occupied with slender tear-shaped and unculturable symbiont fungal cells (Buchner 1965). These described characteristics are also similar to those in Necydalinae. The phylogenetic relationships between Necydalis and these Lepturinae species should be considered in future studies.

The emergence of mycetomes in Spondylidinae and Lepturinae/Necydalinae suggests that the acquisition of this organ is beneficial for these groups. The benefits of this symbiosis need to be considered from the respective perspectives of the symbionts and longicorn beetles. Establishing symbiotic relationships between Cerambycidae through mycetomes would have apparent advantages for fungi. Our study and previous studies have shown that the mycetomes of some species include fungi that cannot proliferate on a PDA plate, although such a plate is sufficient for many yeasts to grow (Müller, 1934). From this result, one of the functions of the mycetome is to provide fungi with nutrients necessary for their proliferation.

Moreover, we here showed that the mycetomes of larval Lepturinae possessed yeast cells even when the larvae ate an artificial diet containing anti-fungal chemicals, which can inactivate yeast cells by overnight incubation, suggesting that another role of the mycetome is to protect fungi from the chemicals that could be included in plants or secreted from competitive fungi. Unlike larvae of the species in Lepturinae, *Necydalis formosana* larvae lost symbionts from their mycetomes when cultured with a diet containing anti-fungal chemicals, suggesting that the mycetomes of Lepturinae and Necydalinae are functionally and structurally different. In most cases, a single yeast species occupies the mycetome of a larva, even though the larvae have a chance to incorporate external yeasts or yeast-like fungi that are spontaneously present in dead trees. Therefore, the mycetome can protect symbiotic fungi from competition with other fungi.

In contrast to the advantages for fungi, the benefits conferred to the longicorn beetles remain obscure. Because the mycetome is a longicorn beetle-derived organ and fungi are not necessary for its formation, symbiosis with fungi is advantageous for longicorn beetles. It has been proposed that symbiont fungi are nutritionally beneficial for host beetles, as in the case of many wood-eating insects and their symbiotic gut microorganisms (Gibson and Hunter, 2010). Our study provided evidence suggesting that symbiosis is related to nutrition and/or feeding habits. All species in Spondylidinae examined in this study and most of the mycetome-possessing species in Lepturinae eat brown rot supwood. This shared food choice would have been made possible by the acquisition of mycetomes. Some Lepturinae species do not have mycetomes, and the loss of this organ occurred over several rounds of speciation (Fig. 2). The loss of the mycetome is also strongly associated with the choice of diet. All mycetome-lacking Lepturinae species eat inner bark or living roots. Inner bark and living roots have different nutritional or organismal conditions from those in rot supwood. For example, inner bark and living plant tissue contain more nutrients than supwood (Schowalter 1992). The change in diet preference may have enabled the Lepturinae species to be free from mycetome-dwelling yeasts due to the improved nutrient uptake. When fed an artificial diet, we also showed that mycetome-possessing Lepturinae larvae can grow without symbiont yeasts. For this experiment, we used an artificial diet based on the silkworm diet, which is much richer in nutrients than dead wood. Therefore, the species having mycetomes do not require symbiont yeasts under such improved nutritional conditions, which paradoxically supports the idea that the benefits from symbiont fungi are associated with nutrients. To confirm more conclusively, we need to compare the growth rate between larvae having mycetomes with or those having mycetomes without yeast when fed a diet having low nutritional value. Our method of obtaining larvae with yeast-free mycetomes will assist in future experiments.

Among the Cerambycidae group, many Lamiinae and Cerambycinae species are known to eat inner bark of fresh dead trees or live trees (Svacha and Lawrence 2014). The species in Lamiinae and Cerambycinae do not have mycetomes, probably because their diets are more nutritive than those of mycetome-possessing groups. Lamiinae is the sister group of Spondylidinae, which is dominant in mycetome-possessing species. Unlike Lamiinae, Spondylidinae longicorn beetles in Japan depend on brown rot supwoods (Table S1). The symbiosis strategy with fungi in these two groups may have differed in their diet choices after the split. We have to note that the tendency of diet choice is not always supported since several mycetome-lacking Prioninae (such as *Psephactus remiger*), Cerambycinae (such as *Thranius variegatus*, *Paraclytus excultus*, *Xylotrechus cuneipennis*) and Lamiinae (such as *Nanohammus rufescens*) species eat strongly rot wood or outer bark. These species may have another way to strengthen their nutritional condition, including establishing symbiosis with another microorganism group, such as bacteria, as reported previously (Grünwald et al., 2010; Reid et al., 2011).

In our experiments, mycetome-lacking larvae of longicorn beetles tended not to have yeast in their gut compared to those in mycetome-possessing larvae (Table 1), suggesting that many of mycetome-lacking species do not transfer yeasts between generations. When yeasts were found in these larvae, the assortment of yeast species usually varied among individuals. A possible explanation is that mycetome-lacking larvae uptake yeasts from their environment, and yeast species that can withstand larval digestive systems proliferate in the gut. Such environment-dependent acquisition of yeast is likely to be the ancestral form of the symbiosis between longicorn beetles and yeasts, as was suggested previously (Douglas, 1989), and the external delivery of yeasts observed in mycetome-possessing species is probably the remnant of this ancestral symbiosis. Because the mycetome-lacking longicorn beetle *A. sinicum* can share a limited number of yeast species between generations, the emergence of mycetomes is not requisite for longicorn beetles to initiate symbiosis with specific yeasts.

Because mycetome-possessing longicorn beetles tend to have a single preferred yeast or fungal species/group as their symbiosis partner, mycetomes could limit the symbiont to a specific species. This role of mycetomes should be in concert with the development of the sac in the female genitalia that stores symbiont yeasts (Schomann, 1937; Buchner, 1965). With the aid of the mechanism of vertical transmission, the mycetomes and symbionts have group-specific features. Larvae of the species in Lepturinae tend to have yeasts culturable on the PDA plate and the yeast cells do not fully occupy the lumen of the lobes of mycetomes. Larvae of the species in Spondylidinae have elongated yeasts unculturable on the PDA plate. Larvae of the species in Necydalinae have unculturable fungal cells that are larger than the yeast cells in Lepturinae. The Necydalinae mycetomes are well-developed whose lumen is fully occupied by the fungal cells. The presence of these group-specific characteristics suggests that some limitations worked to prevent the loss of these characteristics even as speciation occurred within these groups. It is possible that the constraint is associated with group-specific diet choices. These group-specific diet choices include the hardness/softness and degree of moisture of rotten wood, and dependency on specific fungal infestation (Svacha and Lawrence, 2014).

However, our data showed that the vertical transmission of symbionts between generations is imperfect, and larvae sometimes have a different yeast from the symbiont species. These exceptions could be explained by somewhat imperfect manner of yeast transmission between generations. Moreover, this accidental “mismatch” could disturb the phylogenetic relationship between symbiont yeasts and longicorn beetle species. During the speciation of longicorn beetles, the acquisition of symbiont fungi from the environment allowed the diversification of symbiont yeast species that is not always associated with longicorn beetle speciation. We found that the Lepturinae species using Symplocaceae as their host predominantly have the symbiont yeast related to *Candida vrieseae*, even though these Lepturinae species are not monophyletic. Probably the use of the same dead tree promoted the sharing of the yeast species between the beetle species. The mismatch between the phylogenies of host beetle species and symbiont yeasts has also been reported in stag beetles (Kubota et al., 2020), suggesting that this incongruent evolution is shared among Coleopteran groups. Such incongruent evolution was probably permitted because symbiont yeasts have similar physiological characteristics and because the function of yeasts may be to provide inessential dietary supplementation, which may have been dispensable when longicorn beetles switched their food from dead wood to inner bark.

## ACKNOWLEDGMENTS

We are grateful to Drs. Ryutaro Iwata, Rikiya Endo, Takahito Watanabe, Yosuke Degawa and Benjamin Harvey for helpful discussions. We also thank Dr. Ryosuke Yamamoto, Tomoji Mikage, Mitsuru Nonaka, Naho Nakao, Haruto Nakao, Keiichi Sasaki and Sigeo Tsuyuki for helping us with the sample collection.

## AUTHOR CONTRIBUTIONS

YS, NY, KM, and MN conceived and designed the study. YS and MN performed most experiments. NY, and KM performed sampling. JY provided the phylogenetic analyses. TH performed molecular experiments. YS wrote the manuscript with the aid of the other authors.

## SUPPLEMENTARY MATERIALS

Supplementary materials for this article are available online.

---

**Table S1.**
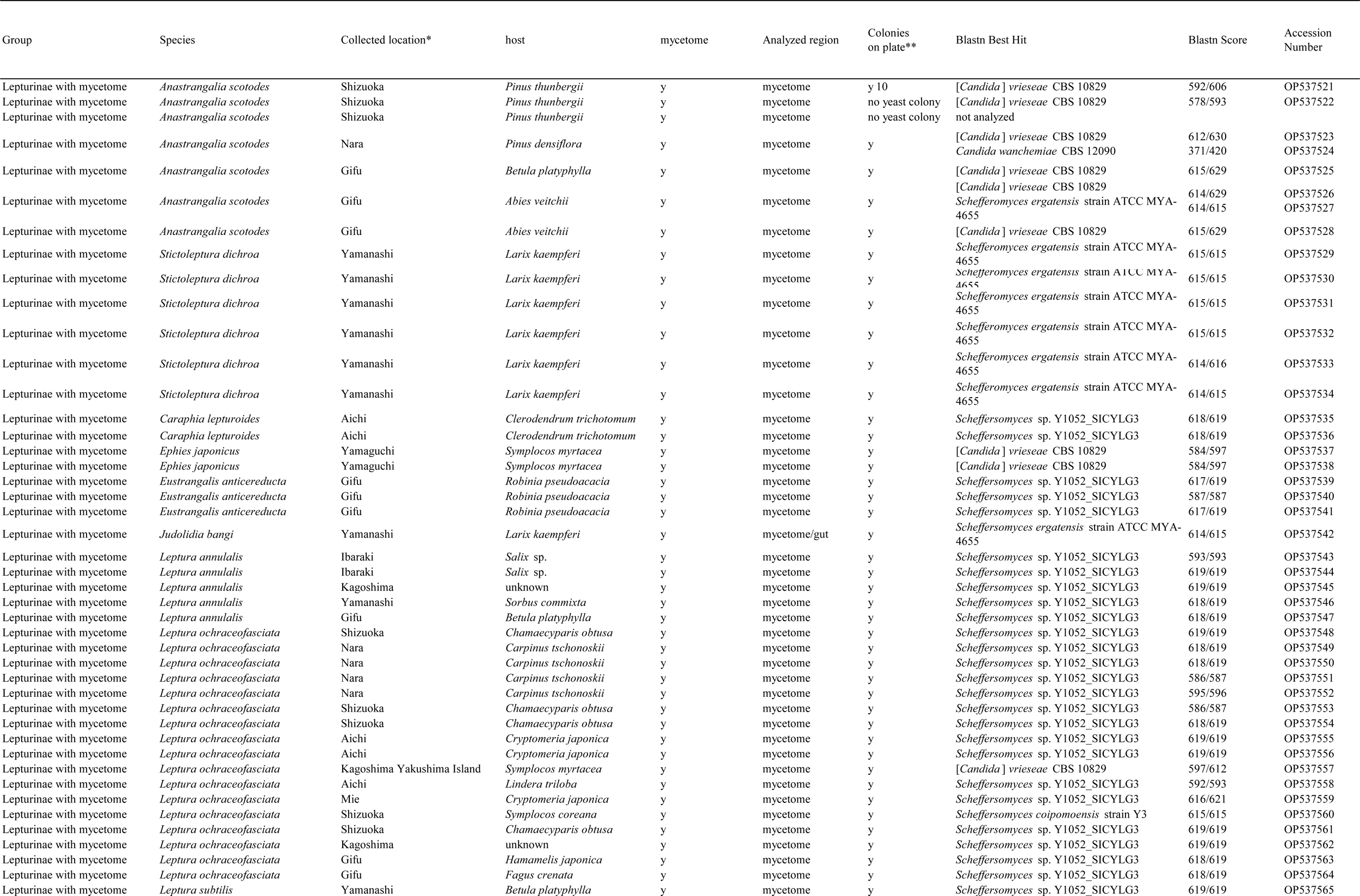
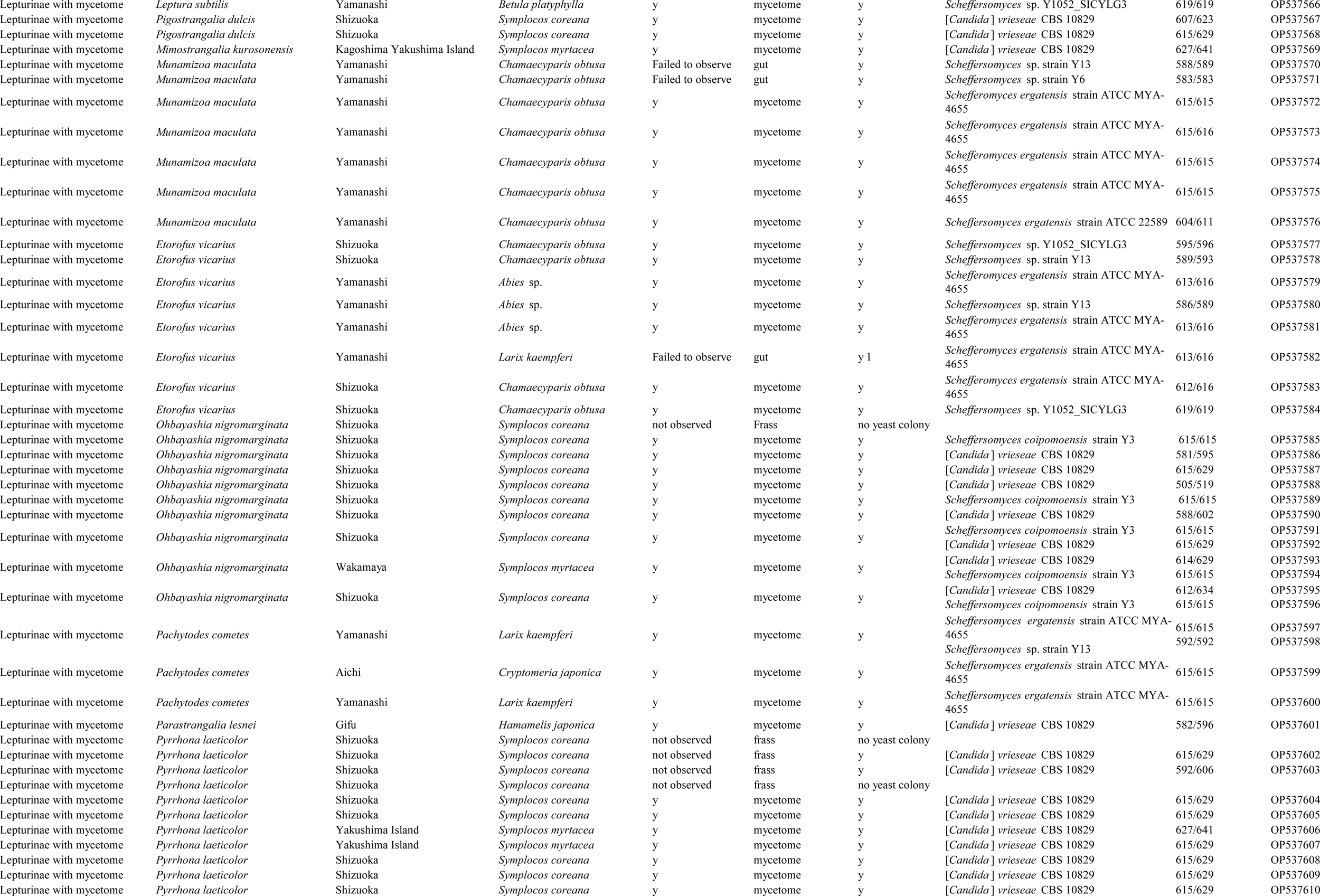
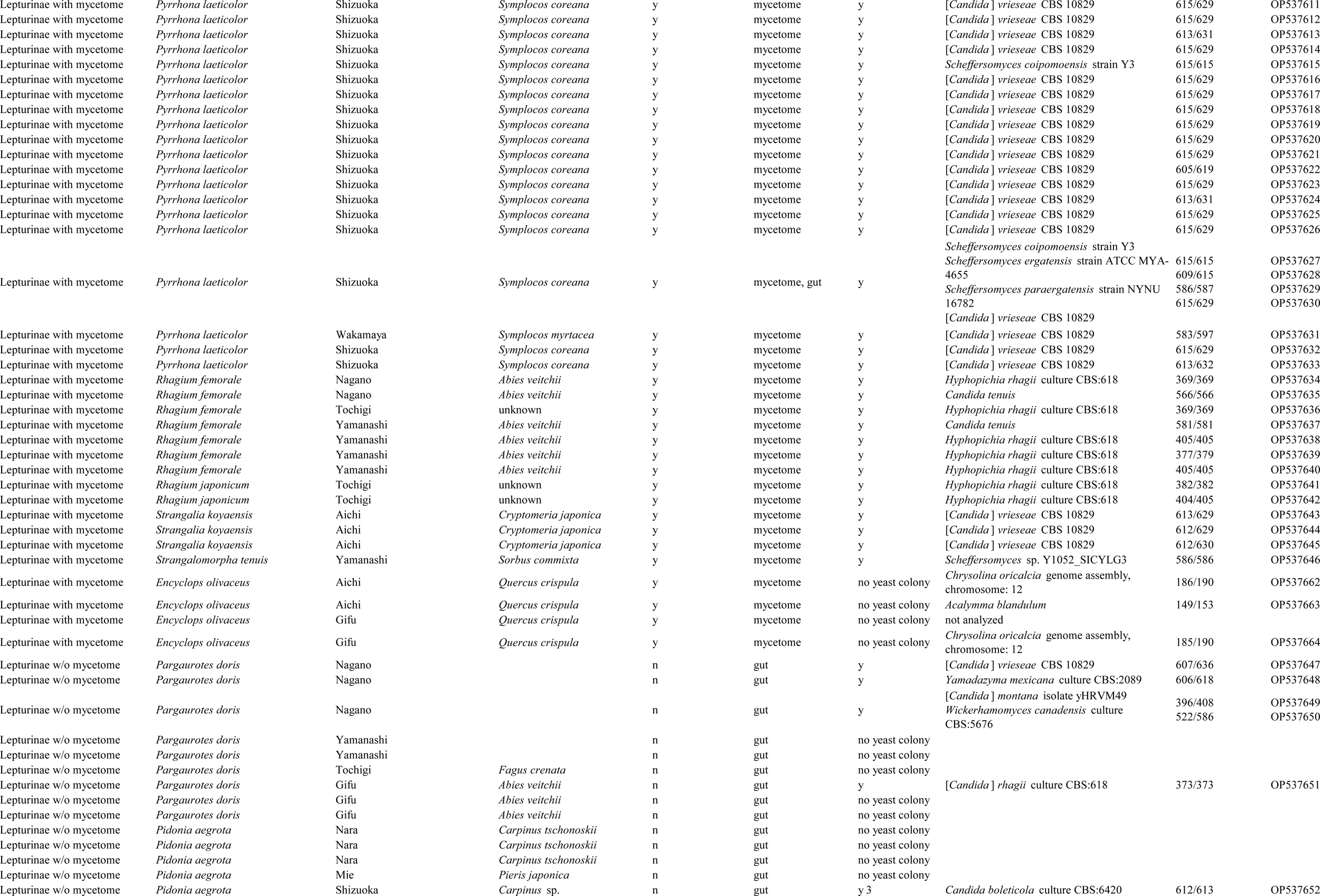
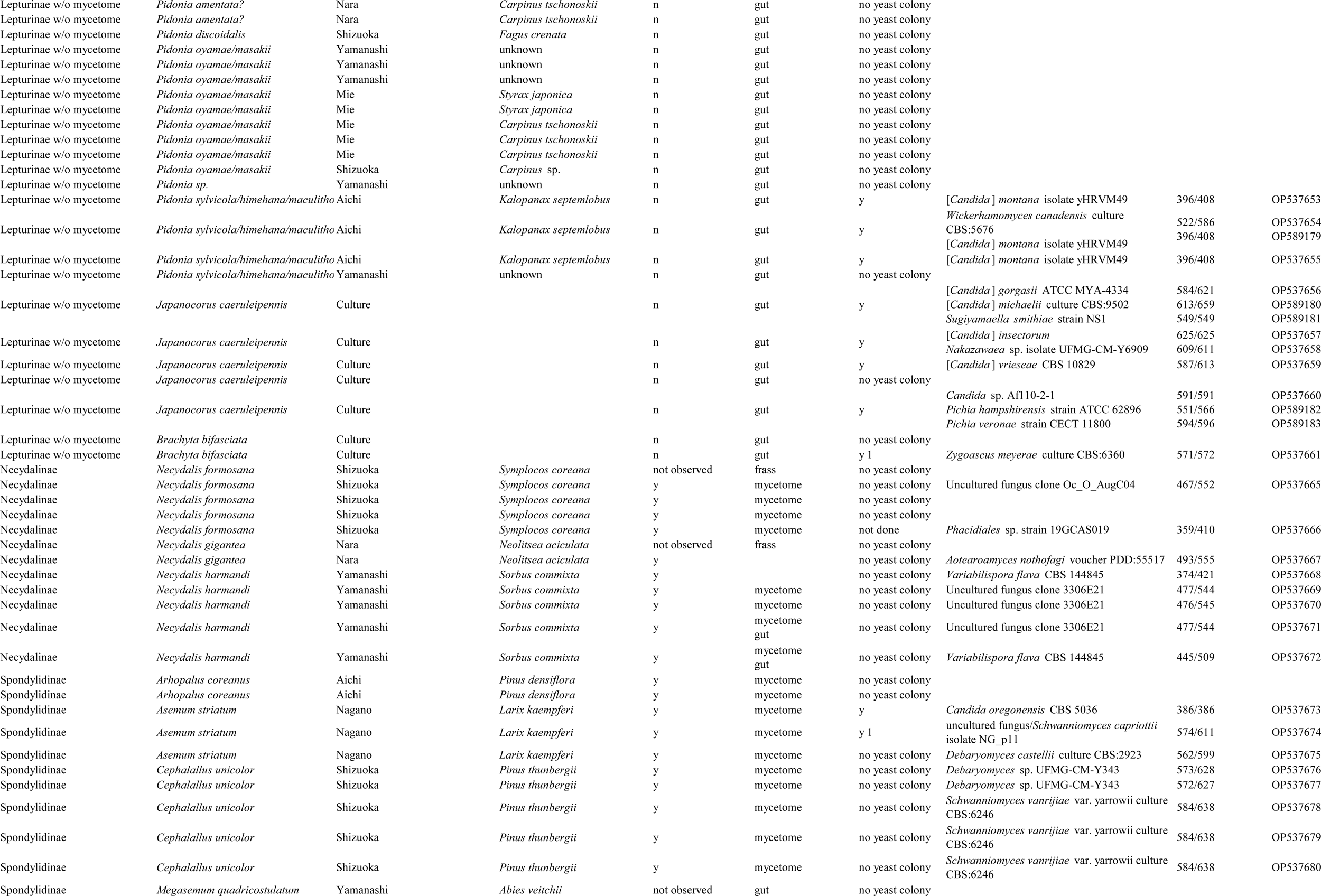
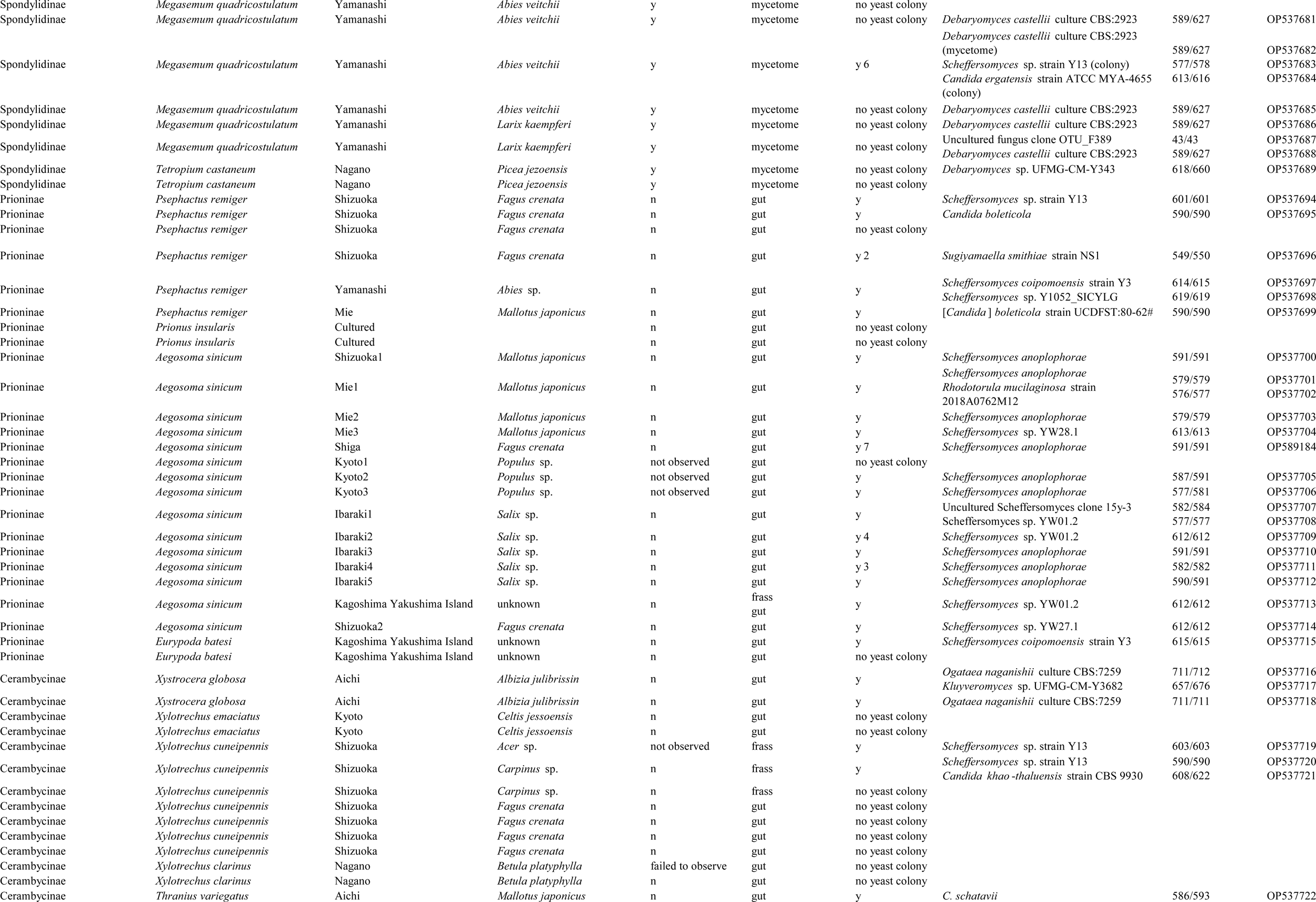
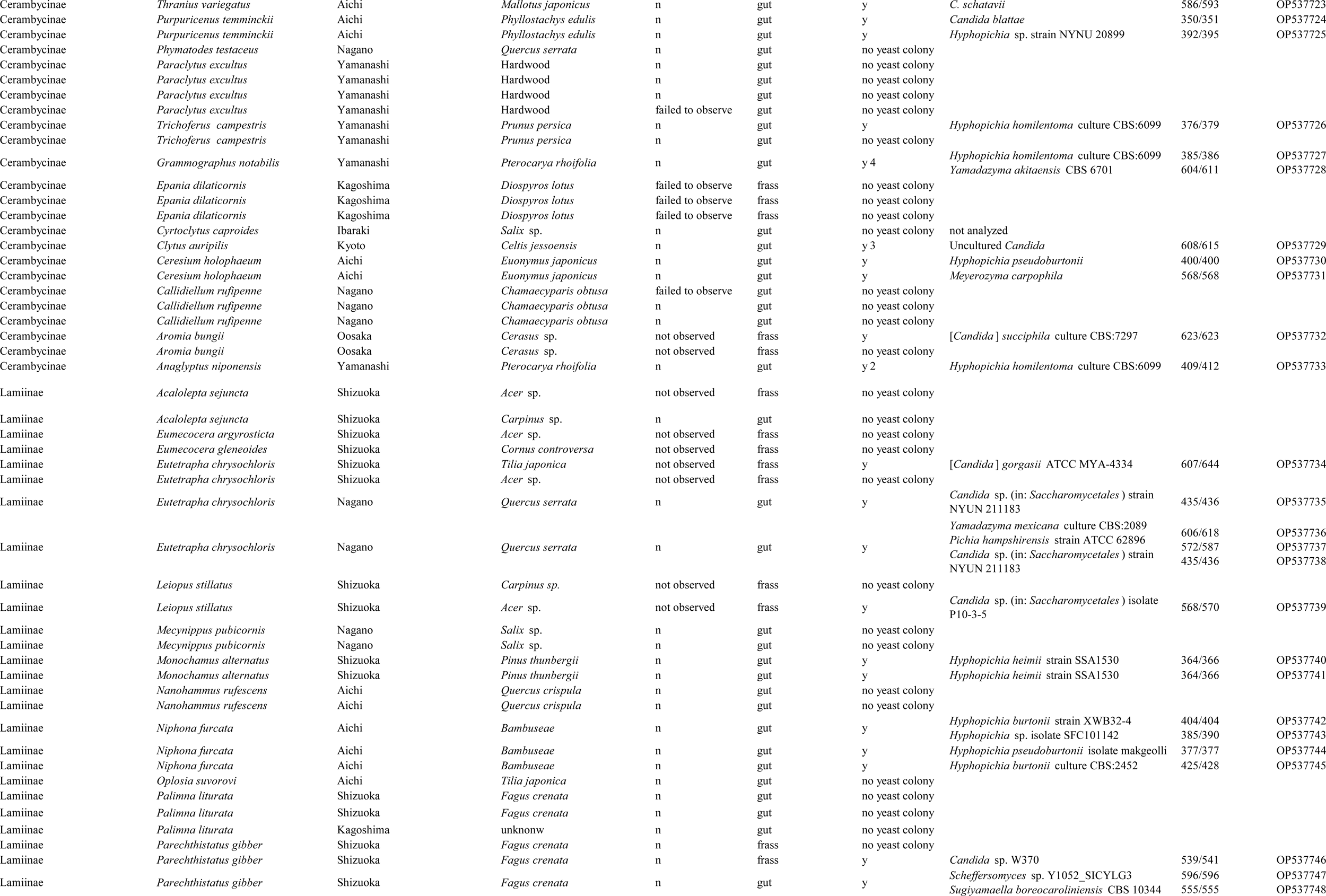
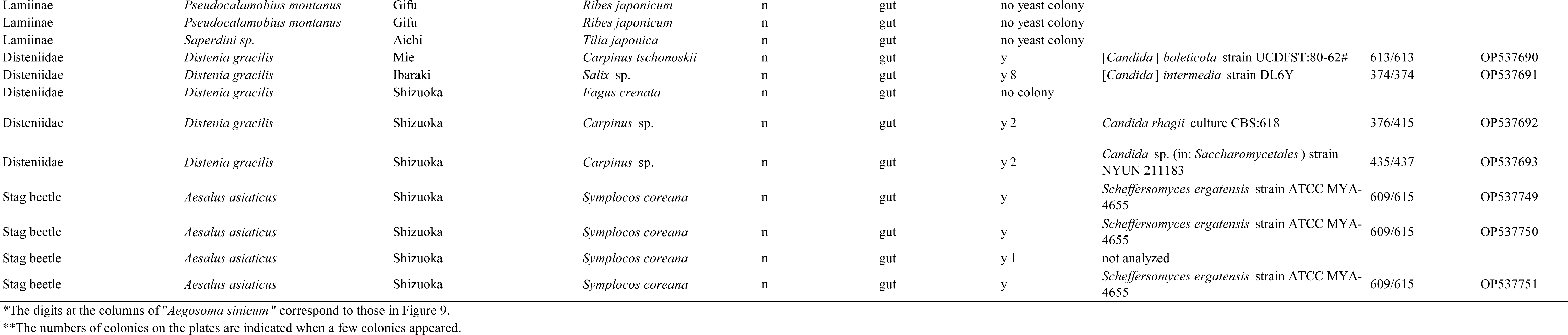
Symbiont fungi isolated from larvae of longicorn beetles in Japan. Each row corresponds to the results of a single larva. “Culture” indicates the cultured larvae used for the analyses. The digits at the columns of “*Aegosoma sinicum*” correspond to those in Figure 9. The numbers of colonies on the plates are indicated when a few colonies appeared.

**Table S2.**
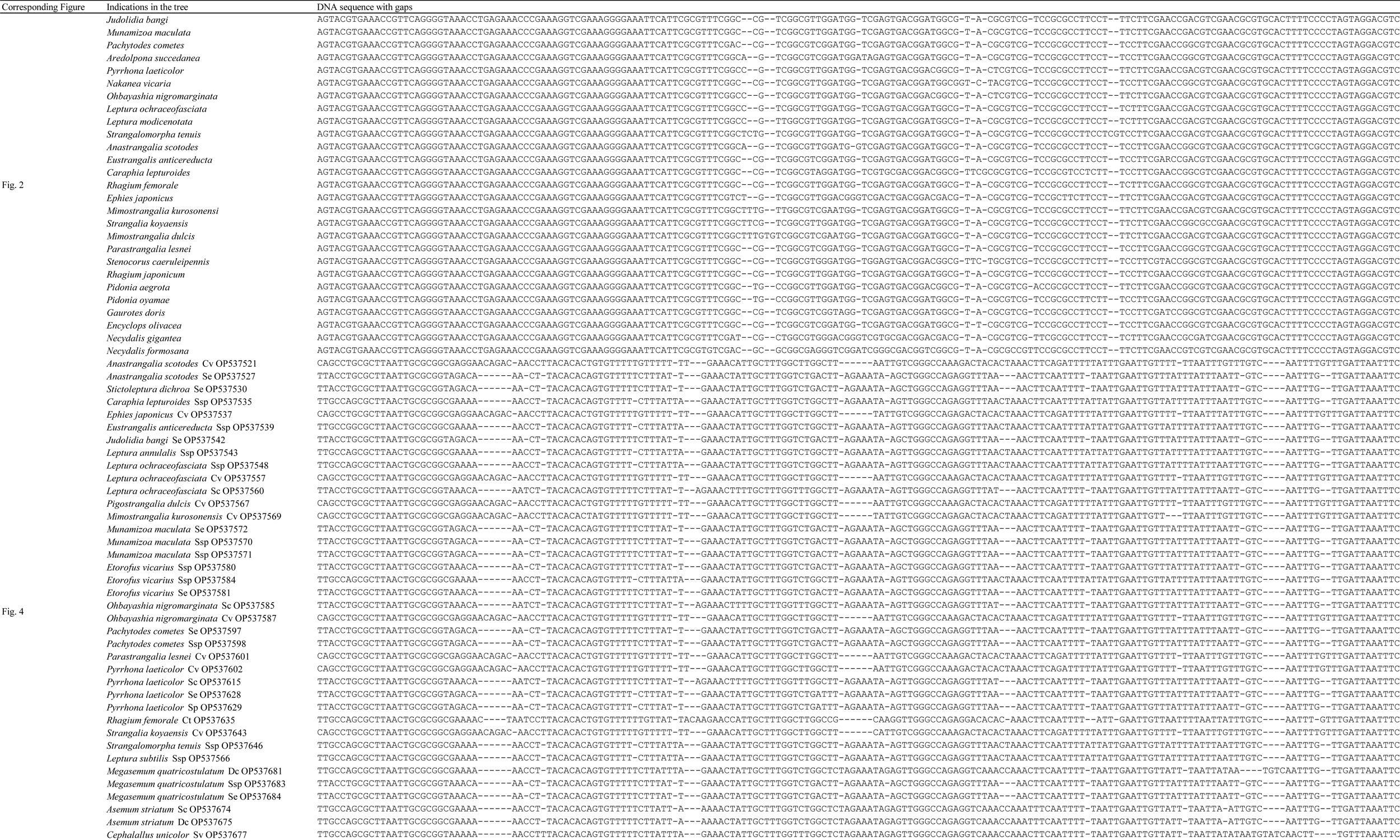
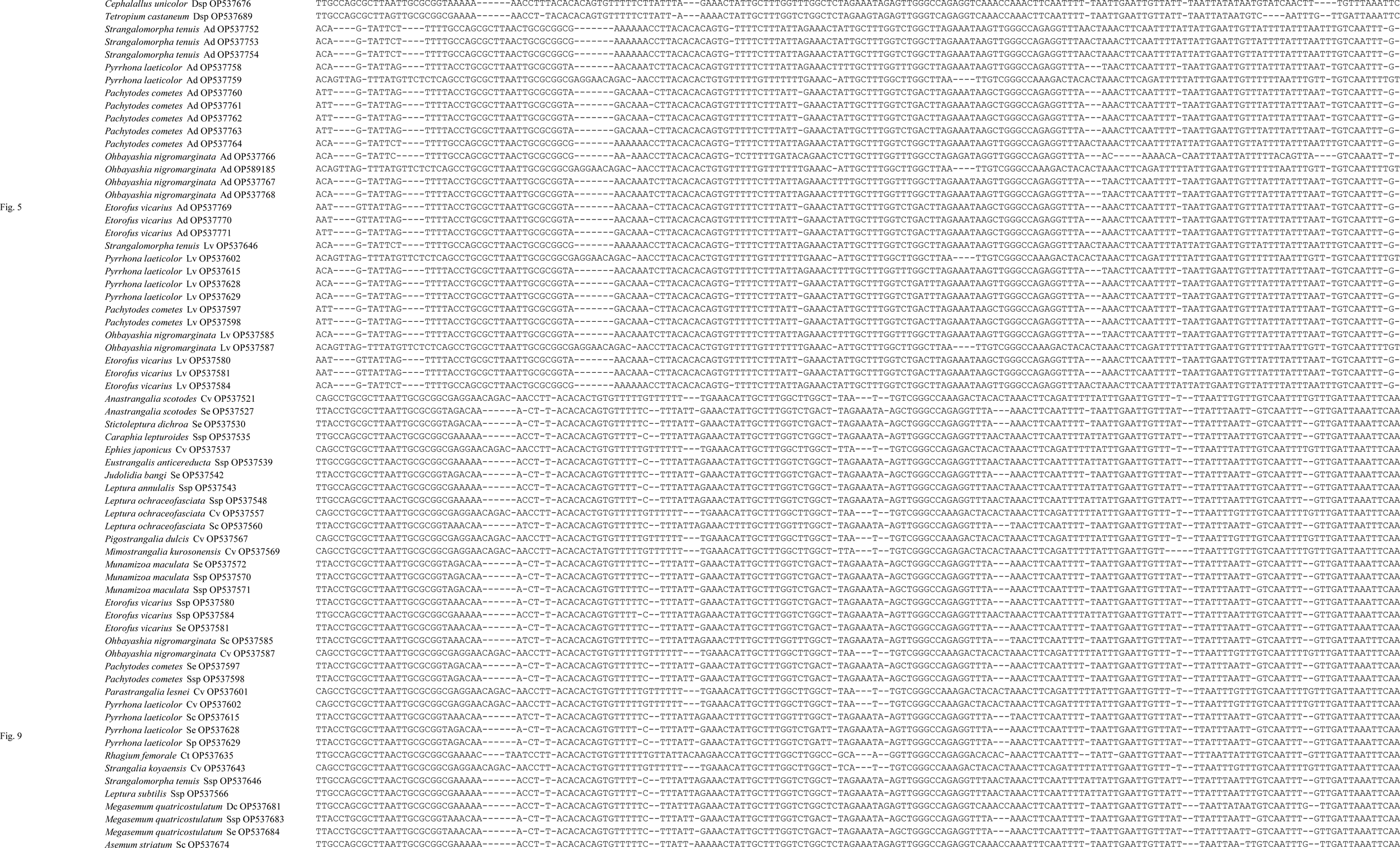
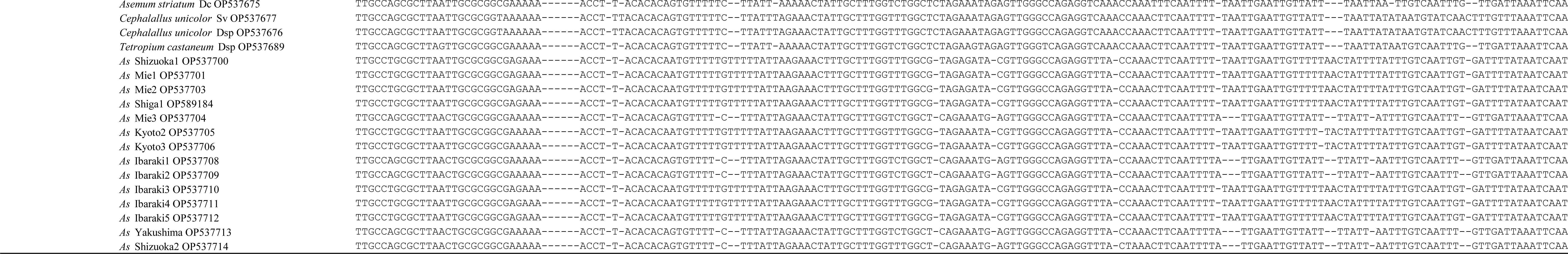
Alignment of nucleotide sequences used for phylogenetic tree construction.

**Table S3.**
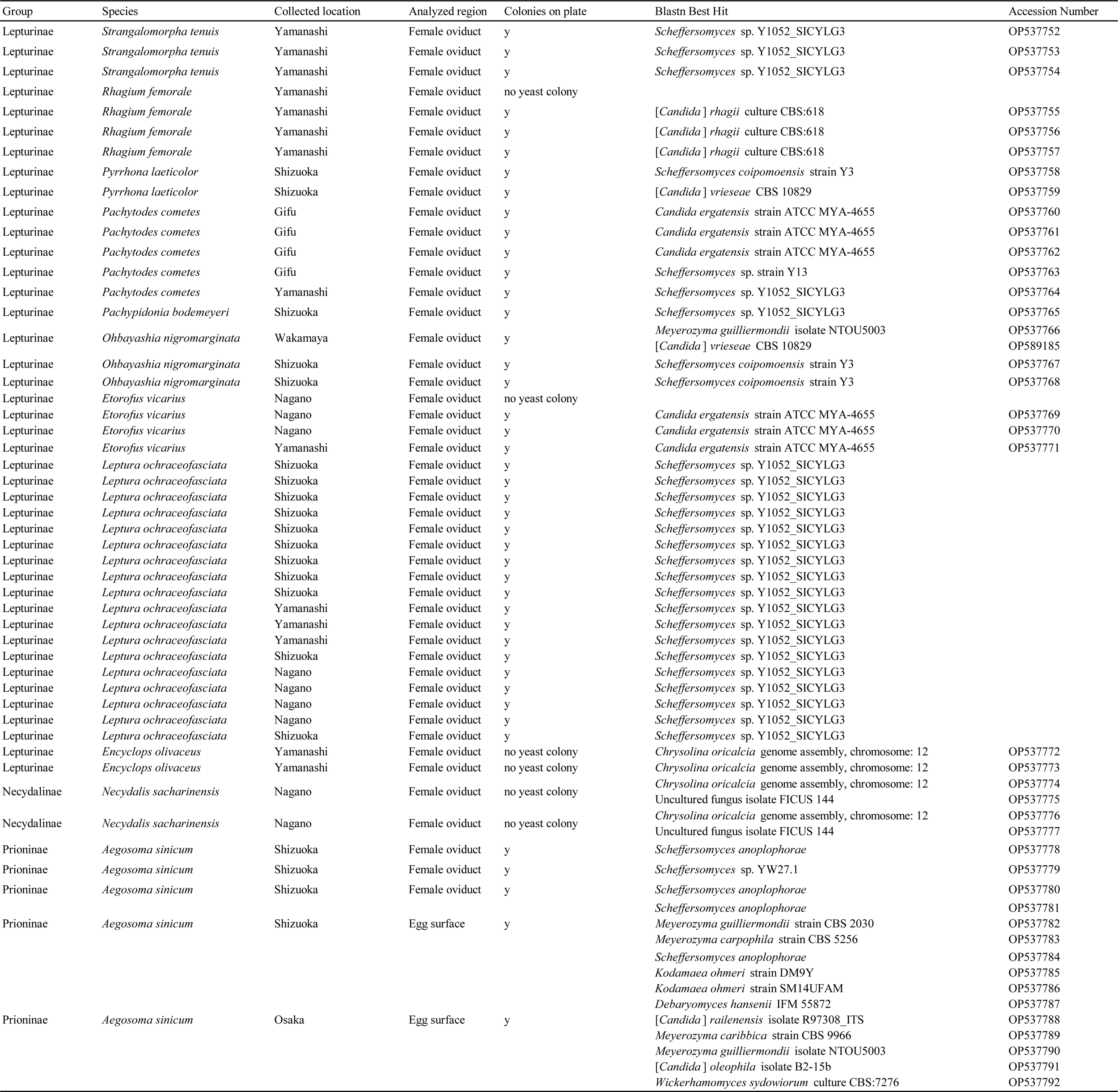
Symbiont fungi isolated from female adults of longicorn beetles in Japan. Each row corresponds to the results of a single female.

**Table S4.**
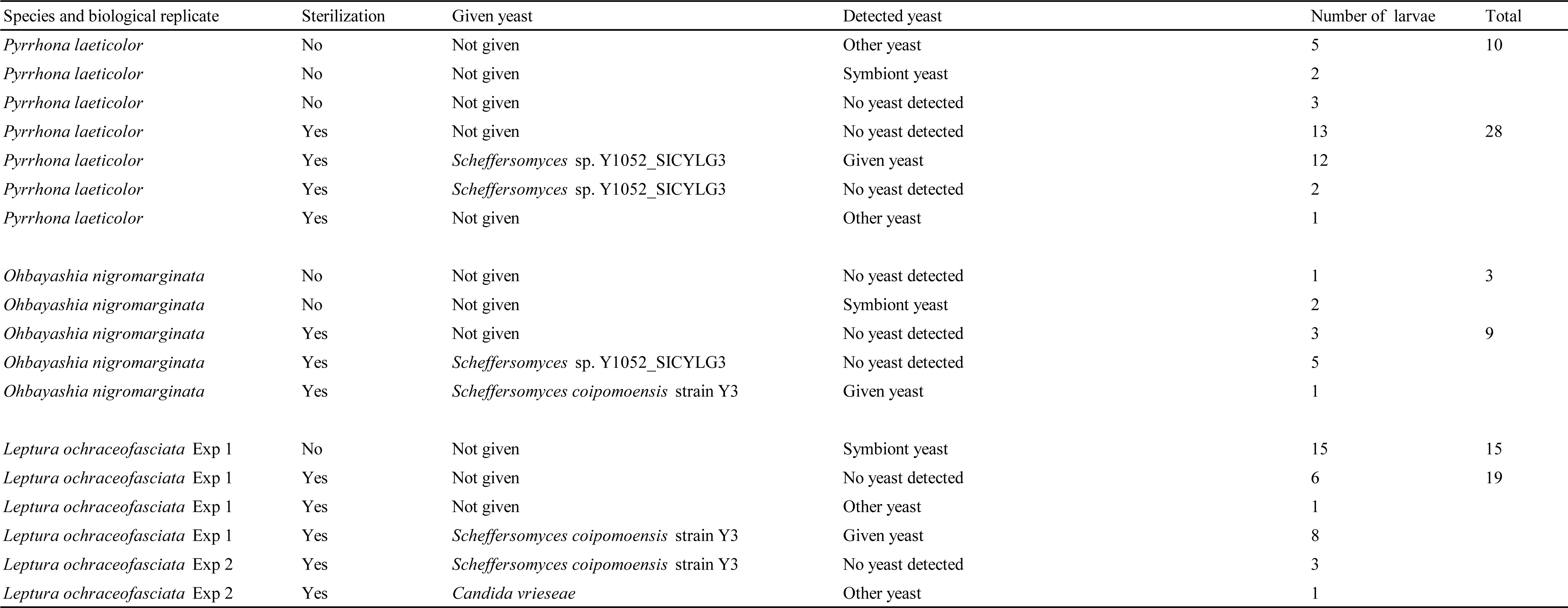
Artificial transfer of yeasts to larvae. The same Exp number indicates the experiment using the eggs from the same female individual.

**Table S5.**
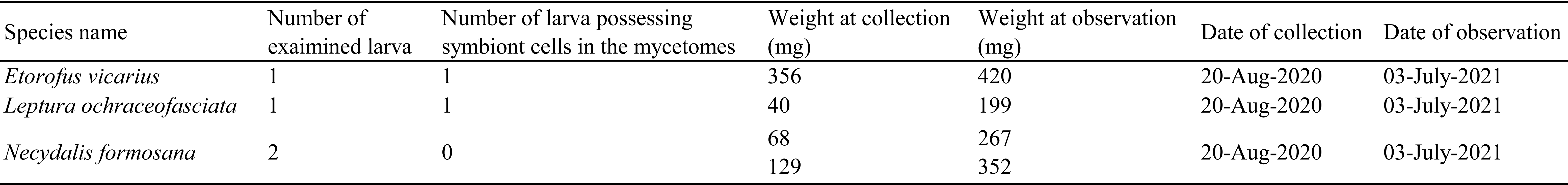
Possession of symbiont cells in the mycetome after long culture in the artificial diet with anti-fungal chemicals.

| Species name | Number of<br>exaimined larva | Number of larva possessing<br>symbiont cells in the mycetomes | Weight at collection<br>(mg) | Weight at observation<br>(mg) | Date of collection | Date of observation |
| --- | --- | --- | --- | --- | --- | --- |
| *Etorofus vicarius* | 1 | 1 | 356 | 420 | 20-Aug-2020 | 03-July-2021 |
| *Leptura ochraceofasciata* | 1 | 1 | 40 | 199 | 20-Aug-2020 | 03-July-2021 |
| *Necydalis formosana* | 2 | 0 | 68<br>129 | 267<br>352 | 20-Aug-2020 | 03-July-2021 |

## REFERENCES

Altschul SF, Madden TL, Schaffer AA, Zhang J, Zhang Z, Miller W, Lipman DJ (1997) Gapped BLAST and PSI-BLAST: a new generation of protein database search programs. Nucleic Acids Res 25: 3389–3402.

Biedermann PHW, Vega FE (2020) Ecology and Evolution of Insect-Fungus Mutualisms. Annu Rev Entomol 65: 431–455.

Buchner P (1965) Symbiosis in insects feeding on cellulose and similar substances. In: Endosymbiosis of animals with plant microorganisms. Interscience Publishers, NY, London, Sydney.

Cherepanov AI (1988) Cerambycidae of Northern Asia. Brill, Leiden, Netherlands.

Danilevsky M (2020) Catalogue of Palaearctic Coleoptera Volume 6/1 (Revised and updated second edition) Chrysomeloidea I (Vesperidae, Disteniidae, Cerambycidae) Brill, Leiden, Netherlands.

Davis TS (2015) The ecology of yeasts in the bark beetle holobiont: a century of research revisited. Microb Ecol 69: 723–732.

Douglas AE (1989) Mycetocyte symbiosis in insects. Biol Rev 64: 409–434.

Endoh R, Suzuki M, Okada G, Takeuchi Y, Futai K (2011) Fungus symbionts colonizing the galleries of the ambrosia beetle Platypus quercivorus. Microb Ecol 62: 106–120.

Gardes M, Bruns TD (1993) ITS primers with enhanced specifity for Basidiomycetes: application to identification of mycorrhizae and rusts. Mol Ecol 2: 113–118.

Gardiner LM (1970) Rearing wood-boring beetles (Cerambycidae) on artificial diet. Can Entomol 102: 113–117.

Gibson CM, Hunter MS (2010) Extraordinarily widespread and fantastically complex: comparative biology of endosymbiotic bacterial and fungal mutualists of insects. Ecol Lett 13: 223–34.

Haddad S, Shin S, Lemmon AR, Lemmon EM, Svacha P, Farrell B, Slipinski A, Windsor D, McKenna DD (2018) Anchored hybrid enrichment provides new insights into the phylogeny and evolution of longhorned beetles (Cerambycidae). Syst Entomol 43: 68–89.

Hulcr J, Stelinski LL (2017) The Ambrosia Symbiosis: From Evolutionary Ecology to Practical Management. Annu Rev Entomol 62: 285–303.

Izumitsu K, Hatoh K, Sumita T, Kitade Y, Morita A, Gafur A, Ohta A, Kawai M, Yamanaka T, Neda H, Ota Y, Tanaka C (2012) Rapid and simple preparation of mushroom DNA directly from colonies and fruiting bodies for PCR. Mycosci 53: 396–401.

Joseph R, Keyhani NO (2021) Fungal mutualisms and pathosystems: life and death in the ambrosia beetle mycangia. Appl Microbiol Biotechnol 105: 3393–3410.

Jurzitza G (1960) Zur systematik einiger Cerambycidensymbionten. Arch Mikrobiol 36: 229–243.

Koiwaya S (2017) Kakedashi tengyuya no hisokana yokubou (1) [The secret desires of a longicorn beetle collector] Gekkan-Mushi 560: 28–36 (in Japanese)

Lindsay EC, Metcalfe NB, Llewellyn MS (2020) The potential role of the gut microbiota in shaping host energetics and metabolic rate. J Anim Ecol 89: 2415–2426.

Modolon F, Barno AR, Villela HDM, Peixoto RS (2020) Ecological and biotechnological importance of secondary metabolites produced by coral-associated bacteria. J Appl Microbiol 129: 1441–1457.

Müller W (1934) Untersuchungen über die Symbiose von Tieren mit Pilzen und Bakterien III Mitteilung: Über die Pilzsymbiose holzfressender Insektenlarven. Archiv für Mikrobiologie 5: 84–147.

Nardon P, Grenier AM (1989) Endocytobiosis in Coleoptera: Biological, biochemical and genetic aspects. Ins Endocymbiosis: Morphology, Physiology, Genetics Evolution (ed. by S. W. & G. G.), pp. 175–216. CRC Press, Boca Raton, FL, USA.

Nie R, Vogler AP, Yang X, Lin M (2020) Higher-level phylogeny of longhorn beetles (Coleoptera: Chrysomeloidea) inferred from mitochondrial genomes. Sys Entomol 46: 56–70.

Notredame C, Higgins DG, Heringa J (2000) T-Coffee: A novel method for fast and accurate multiple sequence alignment. J Mol Biol 302: 205–217.

R Core Team (2021) R: A language and environment for statistical computing. R Foundation for Statistical Computing, Vienna, Austria. URL https://www.R-project.org/.

RStudio Team (2020) RStudio: Integrated Development for R. RStudio, PBC, Boston, MA URL http://www.rstudio.com/.

Schomann H (1937) Die Symbiose der Bockkäfer. Zeitschrift für Morphologie und Ökologie der Tiere 32: 542–612.

Suh SO, McHugh JV, Pollock DD, Blackwell M (2005) The beetle gut: a hyperdiverse source of novel yeasts. Mycol Res 109: 261–265.

Svacha P, Lawrence, JF (2014) In: Handbook of Zoology, Arthropoda. Insecta. Coleoptera. Volume 3: Coleoptera, Beetles. Morphology and Systematics (Phytophaga). Ed by Kristensen NP, Beutel RG, pp 16-177. De Gruyter, Berlin, Germany.

Tanahashi M, Kubota K, Matsushita N, Togashi K (2010) Discovery of mycangia and the associated xylose-fermenting yeasts in stag beetles (Coleoptera: Lucanidae). Naturwissenschaften 97: 311–317.

Tamura K, Stecher G, Kumar S (2021) MEGA11: Molecular Evolutionary Genetics Analysis Version 11. Mol Biol Evol 38: 3022–3027.

Toki W, Takahashi Y, Togashi K (2013) Fungal garden making inside bamboos by a non-social fungus-growing beetle. PLoS One 8: e79515.

Vega FE, Blackwell M (2005) Insect-fungal associations: ecology and evolution. Oxford University Press, Oxford.

Wallace IM, O’Sullivan O, Higgins DG, Notredame C (2006) M-Coffee: combining multiple sequence alignment methods with T-Coffee. Nucleic Acids Res 34: 1692–1699.

White T, Bruns T, Lee S, Taylor J (1990) Amplification and direct sequencing of fungal ribosomal RNA genes for phylogenetics. In: PCR protocols, a guide to methods and applications. Ed by M Innis, D Gelfand, J Sninsky, T White, pp. 315-322. Academic Press, CA, USA.

